# Transcriptome profiling of the *Caenorhabditis elegans* intestine reveals that ELT-2 negatively and positively regulates intestinal gene expression within the context of a gene regulatory network

**DOI:** 10.1101/2021.08.29.457787

**Authors:** Robert T.P. Williams, David C. King, Izabella R. Mastroianni, Jessica L. Hill, Nicolai W. Apenes, Gabriela Ramirez, E. Catherine Miner, Andrew Moore, Karissa Coleman, Erin Osborne Nishimura

## Abstract

ELT-2 is the major transcription factor required for *Caenorhabditis elegans* intestinal development. ELT-2 expression initiates in embryos to promote development and then persists after hatching through the larval and adult stages. Though the sites of ELT-2 binding are characterized and the transcriptional changes that result from ELT-2 depletion are known, an intestine-specific transcriptome profile spanning developmental time has been missing. We generated this dataset by performing Fluorescence Activated Cell Sorting (FACS) on intestine cells at distinct developmental stages. We analyzed this dataset in conjunction with previously conducted ELT-2 studies to evaluate the role of ELT-2 in directing the intestinal gene regulatory network through development. We found that only 33% of intestine-enriched genes in the embryo were direct targets of ELT-2 but that number increased to 75% by the L3 stage. This suggests additional transcription factors promote intestinal transcription especially in the embryo. Furthermore, only half of ELT-2’s direct target genes were dependent on ELT-2 for their proper expression levels, and an equal proportion of those responded to *elt-2* depletion with over-expression as with under-expression. That is, ELT-2 can either activate or repress direct target genes. Additionally, we observed that ELT-2 repressed its own promoter, implicating new models for its autoregulation. Together, our results illustrate that ELT-2 impacts roughly 20 – 50% of intestine-specific genes, that ELT-2 both positively and negatively controls its direct targets, and that the current model of the intestinal regulatory network is incomplete as the factors responsible for directing the expression of many intestinal genes remain unknown.

## Introduction

Transcription factors (TFs) work together to form gene regulatory networks (GRNs) that direct the expression of target genes (Davidson et al. 2003; Levine and Davidson 2005; Emmert-Streib et al. 2014). These GRNs form positive, negative, feedforward, and auto-inhibitory connections to orchestrate downstream transcriptional responses critical for development and health (Alon 2007; Murray et al. 2012; Oestreich and Weinmann 2012; Delás and Briscoe 2020). The *Caenorhabditis elegans* intestine is a model for understanding how GRNs contribute to organogenesis (Maduro and Rothman 2002; McGhee 2007; Dimov and Maduro 2019). In embryos, the intestinal GRN initiates through the combined action of maternally loaded transcription factors and inductive cues. The embryonic intestinal GRN is comprised of successive pairs of GATA transcription factors that culminate in the expression of the GATA transcription factor ELT-2 (McGhee 2007; Maduro 2019). ELT-2 promotes intestine-specific gene expression, its loss leads to a larval lethal intestinal phenotype, and its misexpression ectopically induces intestinal features, all characteristics underscoring ELT-2’s central role (Fukushige et al. 1998; McGhee et al. 2007, 2009; Riddle et al. 2013, 2016; Du et al. 2016; Choi et al. 2017). However, ELT-2 is induced by and co-expressed with other members of the intestinal GRN that together promote cellular specification, commitment, differentiation, and downstream biological processes (Maduro and Rothman 2002; Dimov and Maduro 2019). A missing aspect of the *C. elegans* intestine GRN model is how it changes over developmental time, especially beyond embryonic stages. Many TFs within the described embryonic network are transient, though ELT-2 and its partner ELT-7 remain expressed for the duration of the worm’s lifespan, prompting the questions of whether and how the intestine transcriptional landscape remains consistent or changes over developmental time (Mann et al. 2016; Dineen et al. 2018).

The developmental progression that gives rise to the *C. elegans* intestine is well established. The intestine arises clonally from a single cell, the E cell, and ultimately produces a 20-cell-long tube that extends from the animal’s pharynx to its rectum. At the 4-cell stage, specification of the endomesoderm occurs when a non-canonical Wnt-signaling pathway responds to positional cues to alleviate endoderm repression orchestrated by SKN-1 (Maduro and Rothman 2002; Maduro et al. 2005b; Wiesenfahrt et al. 2015). Beginning in the E-cell (8-cell stage of embryogenesis), transient pulses of GATA TF pairs, first MED-1/MED-2 then END-1/END-3, lead to the eventual expression of the final pair ELT-2 and ELT-7 (Maduro et al. 2005a; Sommermann et al. 2010; Choi et al. 2017; Dimov and Maduro 2019; Maduro 2019; Ewe et al. 2022). ELT-2 and ELT-7 are partially redundant TFs whose expression is sustained through embryonic, larval, and adult stages, only declining in aged, post-fertile worms (McGhee et al. 2009; Sommermann et al. 2010; Mann et al. 2016; Dineen et al. 2018). After initial ELT-2 expression, the intestine continues to divide and grow, undergoing a further 3-4 rounds of cell division. During these divisions, the E lineage ingresses, aligns into two rows of cells down the embryo center, fuses, and finally creates a lumen to form the alimentary canal (McGhee 2007). During larval and adult stages, ELT-2 continues to mark intestinal identity and contributes to diverse intestinal processes such as digestion, immunity, detoxification, and aging (Elliott et al. 2010; Head and Aballay 2014; Roh et al. 2014; Head et al. 2016; Mann et al. 2016; Yang et al. 2016; Su et al. 2020; Zárate-Potes et al. 2020).

Previous analyses identified that the intestine transcriptome is developmentally dynamic as only 20% of genes are shared between embryonic and larval stages as initially assessed by SAGE analysis, a forerunner of microarrays (McGhee et al. 2007, 2009). These changes illustrate the multiple functions the intestine undertakes upon hatching. Because ELT-2 DNA binding sites (TGATAA) are over-represented in the promoters of genes expressed in both embryonic and larval stages, ELT-2 was initially suggested to directly activate all intestinal genes. However, ChIP-seq assays of ELT-2 have revealed that ELT-2 binds only a small fraction of potential binding sites, echoing a general theme in the TF field that sequence alone does not dictate TF occupancy (Carr and Biggin 1999; Liu et al. 2006; Yang et al. 2006; McGhee et al. 2009; Wiesenfahrt et al. 2015; Mann et al. 2016; Kudron et al. 2017). Chromatin, flanking sequences, long-range interactions, and combinatorial TF binding also contribute to the final set of genomic loci a TF inhabits. Indeed, recent evidence suggests a quantitative relationship between ELT-2 cis-regulatory site binding affinity and transcriptional strength (Lancaster and McGhee 2020). However, TFs differ in the degree to which their occupancy at a genomic locus leads to functional transcriptional activation (Ucar et al. 2009). Altogether, differences in intestinal transcriptomics over time could be governed by a host of different molecular changes at the chromatin, DNA, or transcription factor levels.

To date, a systems biology investigation of ELT-2’s regulatory role in the developing intestine utilizing whole genome approaches has yet to be performed. This is largely because the field has lacked a comprehensive transcriptome profile of the intestine over developmental time. To amend this issue, we characterized intestinal transcriptomes in embryo, L1, and L3 stages by Fluorescence Activated Cell Sorting (FACS) followed by bulk RNA-seq. We then combined that dataset with publically available datasets, including ELT-2 DNA binding maps (ChIP-seq) in the same stages, and transcriptional responses to *elt-2* depletion (RNA-seq with or without *elt-2* function at the L1 stage) (Kudron et al. 2017; Dineen et al. 2018).

By integrating the newly generated set of intestine-enriched genes with ELT-2 binding maps and the transcriptional response to *elt-2* deletion, we set out to evaluate three prevailing questions regarding ELT-2 function: 1) Is ELT-2 binding associated with the expression of all intestine genes?, 2) Does ELT-2 function solely as a transcriptional activator? and 3) Does ELT-2 perform positive autoregulation? We found that only 33% of embryo stage intestine-enriched genes had observable ELT-2 occupancy in ChIP-seq assays but that this number increased to 75% in the L3 stage. This suggests that intestinal gene expression in the embryo is supported by additional TFs. Further, our analysis suggests that ELT-2 both positively and negatively regulates subsets of direct target genes. This finding led us to reevaluate the hypothesis that ELT-2 performs positive autoregulation and demonstrated that ELT-2 negatively regulates target genes as well as its own promoter. By using systems biology approaches, we have characterized the complexity of ELT-2’s role in intestinal transcriptional regulation. As more ChIP-seq datasets become available through the modERN Resource (model organism Encyclopedia of Regulatory Networks) (Kudron et al. 2017), the approaches we undertook here may serve as a roadmap to study other TFs that contribute to diverse C. *elegans* GRNs.

## Materials and methods

### *C. elegans* strains and culture conditions

All worm strains were maintained as described (Stiernagle 2006) and cultured at 20°C on NGM plates seeded with OP50 *E. coli* unless otherwise stated. The wild-type strain N2 (Bristol) was used. Transgenic strains used in this study are below:

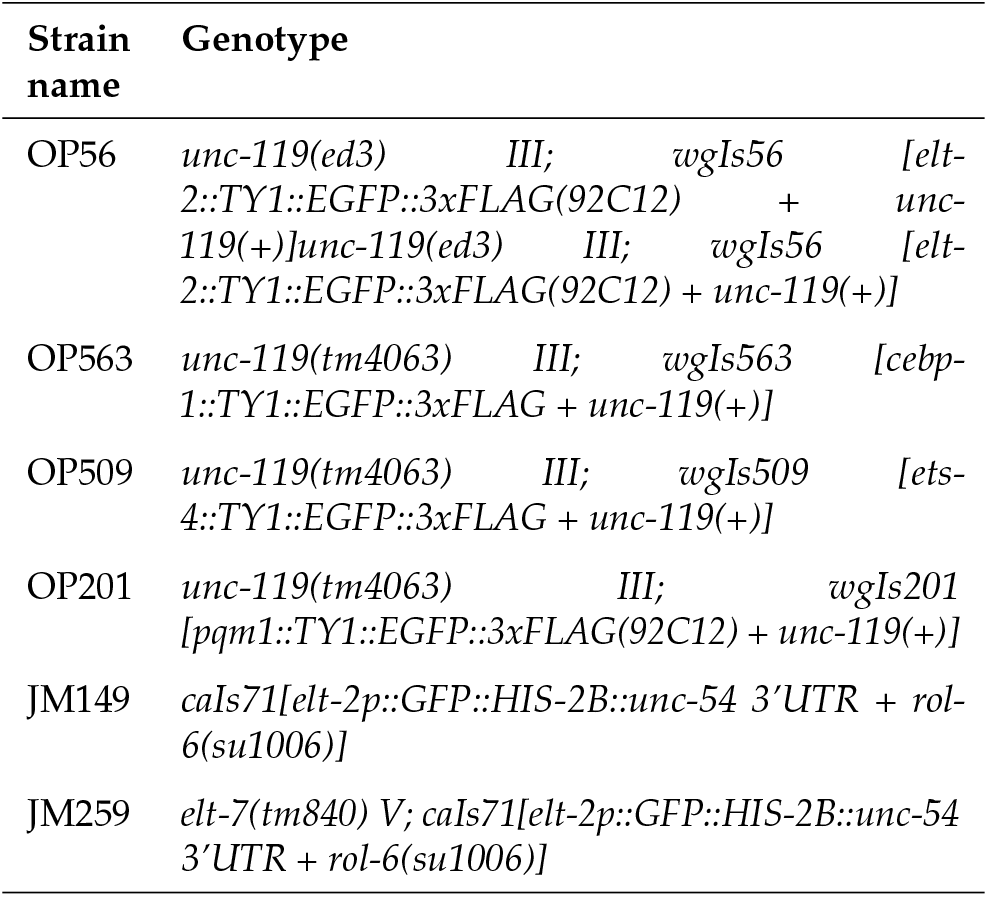

Some strains were provided by the Caenorhabditis Genetics Center, which is funded by NIH Office of Research Infrastructure Programs (P40 OD010440).

### *C. elegans* culture and preparation of dissociated *C. elegans* cells

A detailed protocol for *C. elegans* culture is available on the protocols.io platform: dx.doi.org/10.17504/protocols.io.8epv59zjng1b/ v2. In brief, *C. elegans* were cultured on agar-based NGM plates to reduce any confounding effects that may be introduced by large-scale liquid culture. To produce synchronized cultures sufficient for FACS, two rounds of mixed-stage culture growth were performed followed by two rounds of synchronized growth. *C. elegans* were fed *E. coli* strain OP50 and maintained at 20°C. Synchronized worm populations were achieved through hypochlorite treatment.

For embryo stage experiments, 100,000 synchronized embryos were seeded onto 20 total 150 mm NGM plates at a density of 5,000 embryos per plate. Worms were cultured for 72 hours until gravid and harvested for mixed-stage embryos through hypochlorite treatment. Embryo-stage worms were dissociated through Chitinase and Pronase E treatment and mechanical disruption with a 21-gauge syringe needle. A detailed protocol for embryo dissociation is available on the protocols.io platform: https://dx.doi.org/10.17504/protocols.io.dm6gpbw9plzp/v1

For L1 stage experiments, mixed-stage embryos were synchronized to the L1 stage by incubating in M9 overnight rotating in a 20°C incubator for 24 hours. L1 worms were then fed OP50 E. *coli* on peptone-enriched NGM plates for six hours before cell dissociation to reduce any observable starvation-induced responses in the measured transcriptional data. L1 stage worms were dissociated through SDS-DTT and Pronase E treatment and mechanical disruption with a Dounce homogenizer. A detailed protocol for L1 dissociation is available on the protocols.io platform: https://dx.doi.org/10.17504/protocols.io.rm7vzy365lx1/v1 For L3 stage experiments, synchronized L1 worms were cultured for an additional 48 hours on peptone-enriched NGM plates for 48 hours at 20°C. L3 stage worms were dissociated similarly to L1, where the final centrifugation speed for cell harvest was reduced from 100 rcf to 20 rcf. A detailed protocol for L3 dissociation is available on the protocols.io platform: https://dx.doi.org/10.17504/protocols.io.14egn7zwmv5d/v1.

### FACS isolation of intestine cells

GFP+ intestine cells were isolated from GFP-non-intestine cells by FACS using a BD FACSAria III cell sorter. Additional dyes were used depending on the developmental stage. For embryo stage experiments, viability dye propidium iodide was used to separate live cells (PI-) from dead cells (PI+). For L1 stage experiments, viability dyes were not used because preliminary experiments identified that intestine cells preferentially take up viability dyes confounded sorting (data not shown). For L3 stage experiments, viability dyes are also not used. We observed that L3 stage cell preps have a high degree of debris, so a cell-permeable nucleic acid dye such as DRAQ5 was used to distinguish cells (DRAQ5+) from debris (DRAQ5-). FACS plots and post-hoc analysis were performed with FlowJo (v 10) (Figure 1, S1). Cell staining protocols and gating strategies (Figure S1) are available on the protocols.io platform: https://dx.doi.org/10.17504/protocols.io.j8nlkk43dl5r/v1.

**Figure 1.**
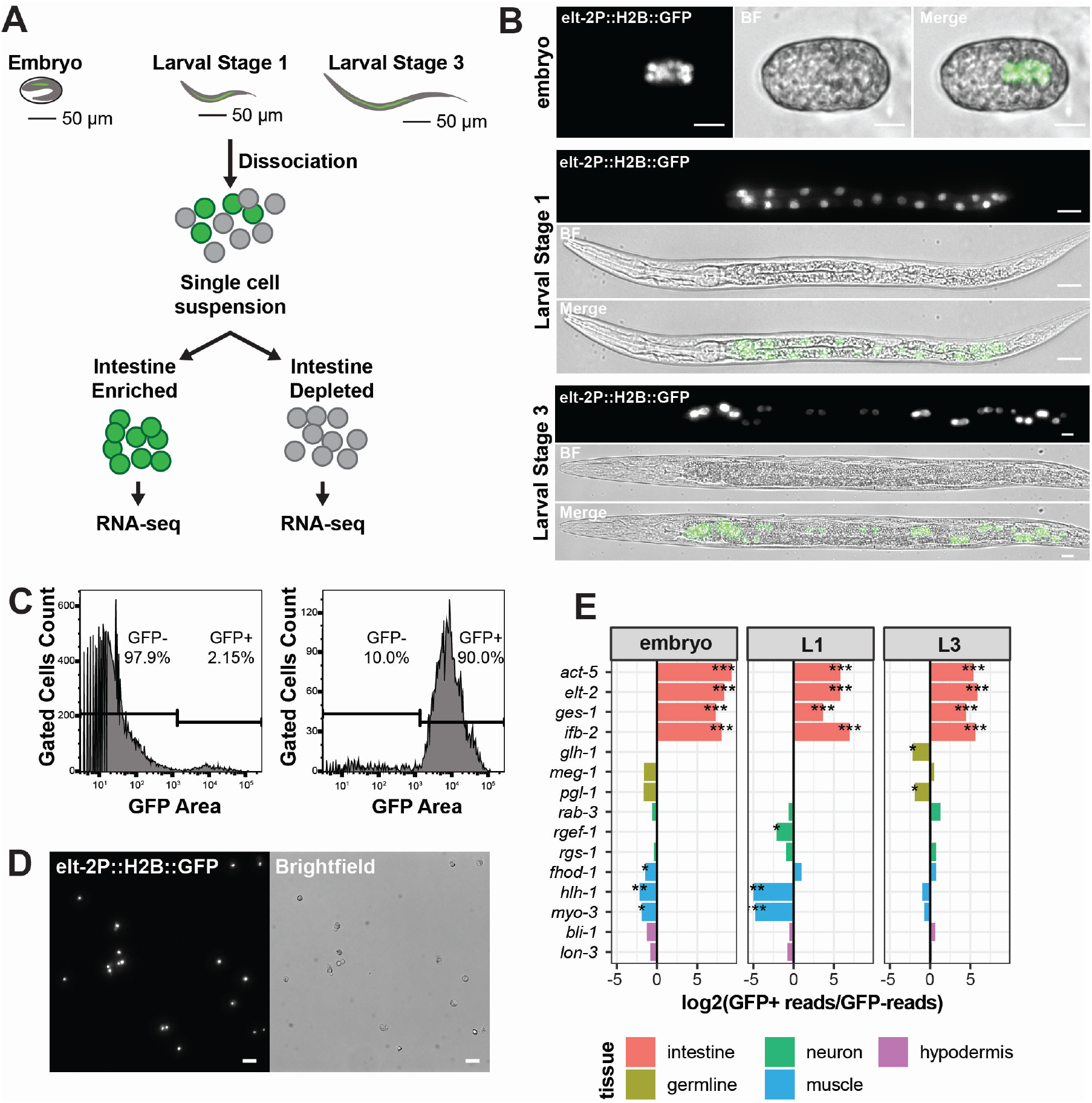
Transcriptional profiling of *C. elegans* intestines using FACS isolation and RNA-seq. (A) Embryonic, L1, or L3 stage worms transgenically expressing H2B::GFP (JM149) in intestinal cells were dissociated into single-cell suspension. Intestine cells were isolated by FACS by GFP fluorescence and profiled using RNA-seq (B) The intestine reporter transgene (*cals71[elt-2p::GFP::HIS-2B]* allele) fluoresces specifically in intestinal cells during embryo (top), L1 (middle), and L3 (bottom) stages (60x magnification, scale bars = 20 microns). (C) Histograms of JM149 dissociated *C. elegans* cells before (left) and after (right) FACS purification. GFP signal gates were set using wildtype strain N2. Percentages of GFP+ and GFP-cells are shown (n > 5,000). (D) Microscopic visualization of GFP+ intestine cells prepared by dissociation and FACS isolation (20x magnification, scale bars = 20 microns). (E) Expression levels of key tissue-specific marker genes in GFP+ (intestine) vs GFP- (non-intestine) samples by RNA-seq (log2 transformed RNA-seq read fold change for tissue-specific genes. The tissue types associated with each marker gene are shown in the color-marked legend.) *P-value < 0.01, **P-value < 1×10^-5^, ***P-value < 1×10^-10^.

Approximately 300,000 GFP+ intestine cells and 1,000,000 GFP-non-intestine cells were collected for RNA isolation for each developmental stage. Three biological replicates for each developmental stage were performed (File S1). FACS isolated cells were pelleted by centrifuging at 10,000 rcf for 5 mins in a 4°C cooled centrifuge. The supernatant was removed from the cell pellet and resuspended in 1 ml Qiazol for RNA extraction. Cells in Qiazol were stored at −80°C until RNA extraction was performed.

### RNA extraction

Total RNA extractions were performed with miRNeasy Micro kit (Qiagen 217084). RNA quantity was measured using a Qubit 3.0 fluorometer (Thermo Fisher Q33216) and Qubit High Sensitivity RNA assay kit (Thermo Fisher Q32852). RNA was of a consistently high quality (RIN > 8.0) as determined on an Agilent 2200 TapeStation using high sensitivity RNA reagents (Agilent 5067-5579). Total RNA samples were stored at −80°C until RNA-seq library preparation was performed.

### RNA-seq library preparation and analysis

To measure the intestine-specific transcriptome, RNA-seq library preparation was performed with the NEBNext Ultra II RNA Library Prep Kit and sequenced with the Illumina platform (NEB E7770S). Ribosomal RNAs (rRNAs) were depleted through the NEBNext Poly(A) mRNA Magnetic Isolation Module (NEB E7490S). All samples were sequenced on the Illumina NovaSeq 6000 instrument except for replicates 1 and 2 for L1 stage samples, which were sequenced on an Illumina HiSeq X-ten instrument. All raw sequencing and processed files are publicly available on NCBI GEO under Accession GSE214581 and accessible at https://www.ncbi.nlm.nih.gov/geo/query/acc.cgi?&acc=GSE214581. The gtf alignment files used in this analysis are also deposited there. The raw sequencing files are available at SRA (Short Read Archive) under Project PRJNA885900.

Sequencing reads were processed using a custom pipeline on the Rocky Mountain Advanced Computing Consortium Summit Supercomputer (File S2). The processing pipeline included the following steps: 1) sequencing reads were filtered for low quality or adapter sequence containing reads with fastp (v 0.20.0) (Chen et al. 2018), 2) aligned to the ce11 *C. elegans* genome (c_elegans.PRJNA13758.WS263.canonical_geneset.gtf) with hisat2 (v 2.1.0) (Kim et al. 2019), 3) tabulated for the number of mapped reads aligning to the WS263 genome assembly using featureCounts (v 1.6.4) (Liao et al. 2014).

Differential expression analysis and plotting were performed with the DESeq2 package (v 1.28.1) in the R Environment (v 4.0.3, BiocManager, v 1.30.10, tidyverse v 1.3.1; File S3) (Love et al. 2014; Wickham et al. 2019; Morgan 2021; Team 2021). RNA-seq count data were filtered for detected genes (> 10 counts per million across all samples). To visualize genome-wide similarities between samples, sample-to-sample Euclidian distances were calculated using the Complete cluster method on normalized, variance-stabilized, log transformed count data (Figure S2). In all pairwise differential expression analyses, genes were identified as differentially expressed if they had a normalized log2-transformed fold change value greater than 1 with a Benjamini-Hochberg corrected p-value less than 0.01 (Figure 2, Figure S3). UpSet plots were generated with the UpSet function of ComplexHeatmap (v 2.8.0) (Gu et al. 2016).

**Figure 2.**
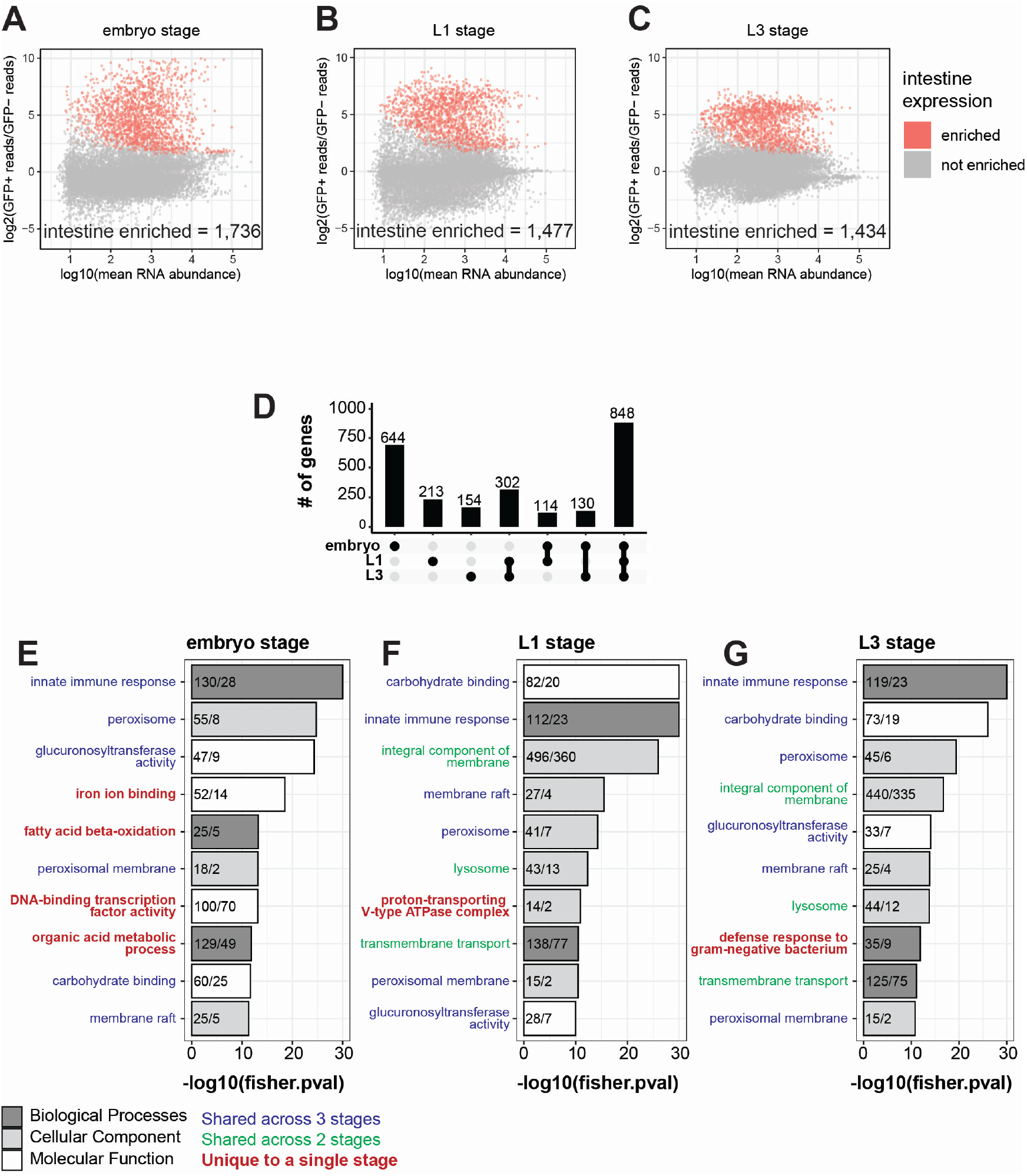
Differential expression analysis identifies intestine-specific genes in embryo, L1 and L3 stages. (A-C) MA-plots illustrate pairwise differential expression in FACS isolated intestine (GFP+) versus non-intestine (GFP-) cells. The log2-fold change of gene expression (y-axis) is plotted against the mean normalized read count per gene (x-axis). Significantly enriched intestine genes are highlighted in red (log2(fold change) > 1 and Benjamini-Hochberg adjusted p-value < 0.01). This analysis identified 1,736 embryo-enriched genes (A), 1,477 L1 intestine-enriched genes, and 1,434 L3 intestine-enriched genes. (D) Upset plot illustrating how the degree of overlap between intestine-enriched sets of genes at different developmental stages. Dots below the x-axis indicate gene set identification. Single dots are specific to one stage while dots connected by lines indicate sets of genes that are shared between stages. (E-G) Gene Ontology (GO) analysis of intestine-enriched genes identified in each developmental stage. The top 10 GO terms for all three categories are displayed (BP, biological process; CC, cellular component; MF, molecular function). The x-axis corresponds to the −log10 transformed p-value, and the y-axis corresponds to the identified GO term. Numbers within each bar represent the number of “observed vs expected” genes in the input set that corresponds to the given GO term. Categorical terms are colored depending on whether they are unique or shared across developmental stages.

### ELT-2 ChIP-seq data and analysis

ELT-2 ChIP-seq data was downloaded from the ModERN Resource for the embryo, L1, and L3 stages (https://epic.gs.washington.edu/modERN/) (Kudron et al. 2017). Downloaded files included aligned reads and optimal IDR thresholded peaks (File S4). Peaks were assigned to genes based on the presence of a ≥ IDR passing peak in a window of −1 kb and +0.2 kb centered on a gene TSS. ELT-2 ChIP-seq heatmaps were generated with deeptools (v 3.5.0) (Ramírez et al. 2016).

### *elt-2(-)* RNA-seq data and analysis

RNA-seq data measuring the transcriptional response to *elt-2* deletion was downloaded from the publication supplemental material (Dineen et al. 2018). Files contained per gene aligned RNA-seq read count data. Differential expression analysis and plotting were performed with the DESeq2 package (v 1.28.1) in the R Environment (v 4.0.3, BiocManager, v 1.30.10, tidyverse v 1.3.1). (Love et al. 2014; Wickham et al. 2019; Morgan 2021; Team 2021). Genes were identified as differentially expressed if they had a normalized log2-transformed fold change value greater than 1 with a Benjamini-Hochberg corrected p-value less than 0.01.

### Gene set definitions

Tissue-specific genes were generated by downloading gene lists associated with the following anatomy ontology terms from WormBase:

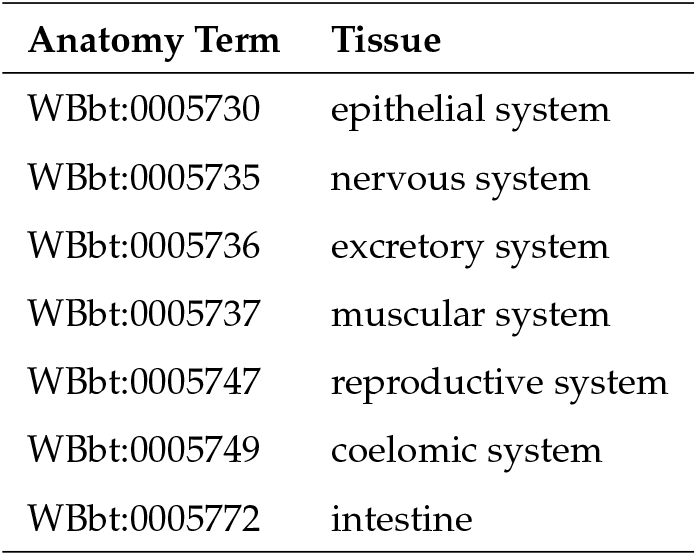

The list of ubiquitous genes was previously defined (Rechtsteiner et al. 2010). Tissue-specific or ubiquitous gene set definitions were made by retaining genes from the input lists that were unique to one category (File S6). The hypergeometric test was utilized to evaluate if a given set of genes corresponded to any gene set listed above. The list of known and putative *C. elegans* transcription factors was provided in the wTF3.0 TF list (Bass et al. 2016).

### Computational Analysis

RNA-seq data and ELT-2 target genes were subset for protein coding genes in all analyses. All analysis and plots were performed in the R Environment unless otherwise stated (v 4.0.3, BiocManager, v 1.30.10, tidyverse v 1.3.1). (Love et al. 2014; Wickham et al. 2019; Morgan 2021; Team 2021). All scripts for analysis are available in the following GitHub repository: https://github.com/rtpwilliams/williams_et_al.

### Gene ontology analysis

Gene Ontology (GO) terms for gene sets were evaluated using R package topGO (v 2.44.0) (Alexa and Rahnenfuhrer 2021). GO analysis was performed using the “weight01” algorithm and the “fisher” statistic. When measuring ontology terms for intestine-enriched FACS samples (Figure 2), background gene sets included all genes in the genome. When measuring ontology terms for ELT-2 regulated targets (Figure 6), background gene sets included all genes with an ELT-2 bound promoter.

**Figure 3.**
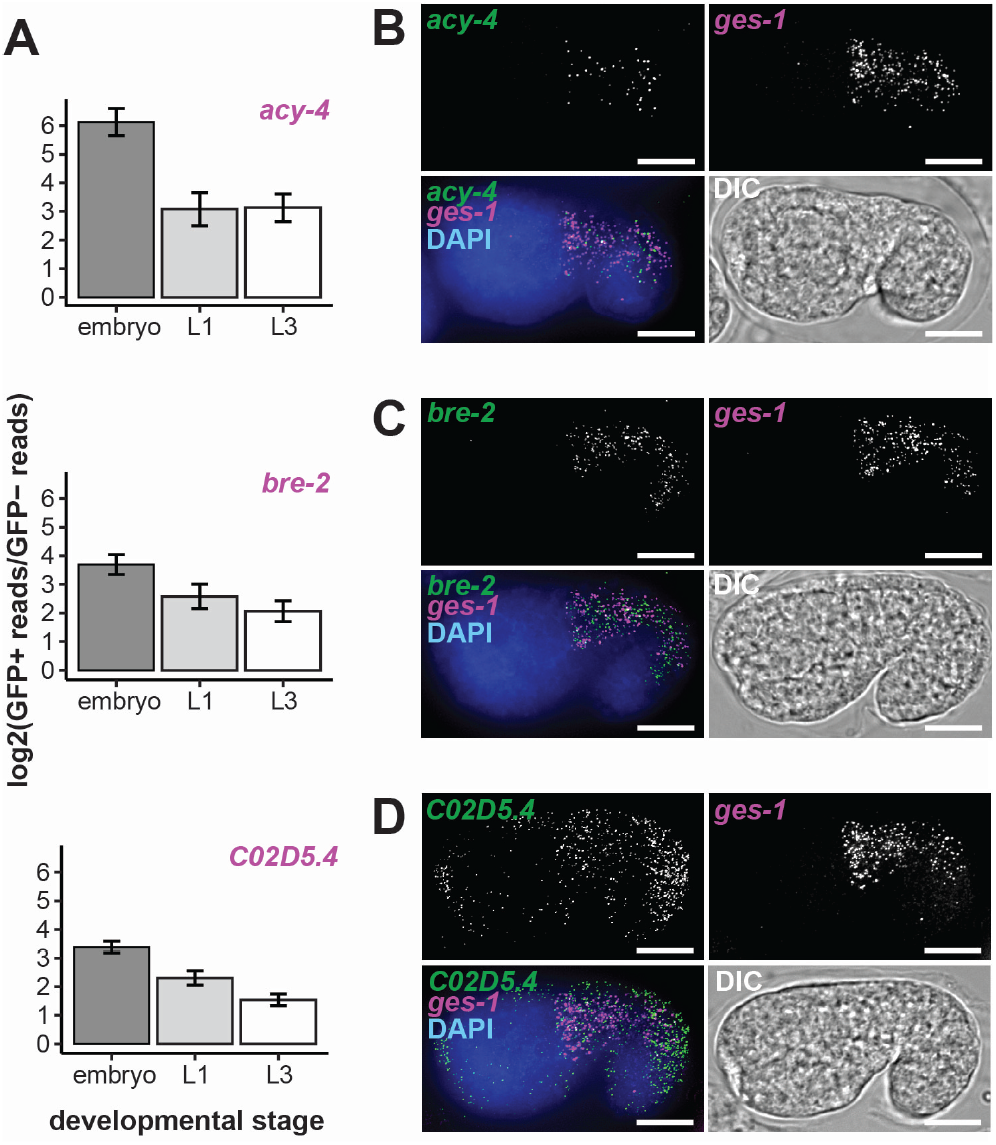
Transcription profiling of FACS sorted intestinal cells identifies new intestine expressed genes. (A) RNA-seq-based intestine-enrichment metrics for *acy-4, bre-2*, C02D5.4 (GFP+ (intestine) vs GFP- (non-intestine) log2 transformed RNA-seq read fold change)). All genes had a log2(fold change) > 1 with Benjamini-Hochberg adjusted p-value < 0.01. Bar plots represent the mean and standard deviation of replicate log2 (fold change) values (embryo: n =3; L1: n=2; L3: n=3)). (B-D) smiFISH (single molecule inexpensive Fluorescence In Situ Hybridization) microscopy images of *acy-4* (B), *bre-2* (C), and C02D5.4 (D) mRNA in comma-stage embryos. Transcript of interest (green), intestine marker transcript ges-1 (magenta), DNA (blue), and DIC images (grey) are shown. Representative images from three biological replicates collected for a total of at least 30 images. Scale bars = 10 microns.

**Figure 4.**
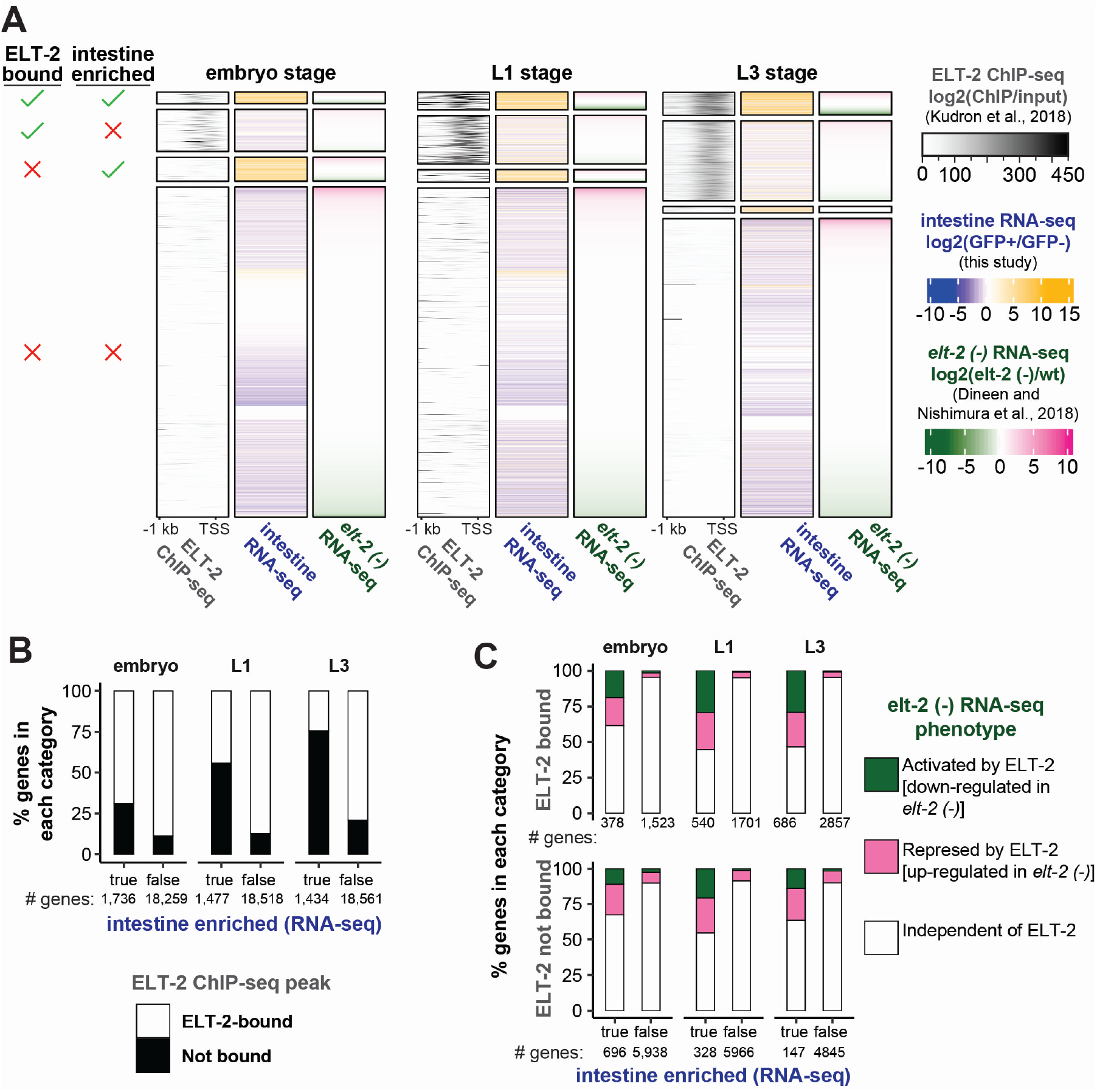
Integration of genome-wide datasets: ELT-2 ChIP-seq, intestine-specific RNA-seq, and elt-2-depleted RNA-seq. (A) Aligned heatmap visualizations of ELT-2 ChIP-seq occupancy (log2 transformed ratio of ChIP-seq/input colored from white to black), intestine-specific RNA-seq fold-change (log2 transformed intestine enrichment colored from-blue to yellow), and *elt-2* depletion transcriptional response (log2 transformed fold change *elt-2(-)* / wt colored from green to pink). ELT-2 binding (ChIP-seq, (Kudron et al. 2017)) and intestine enrichment (RNA-seq, this study) were assessed in embryo, L1 and L3 stages. *elt-2 (-)* transcriptional response (RNA-seq, (Dineen et al. 2018)) was measured in the L1 stage. Gene sets were separated based on ELT-2 binding and intestine enriched expression. Genes within each category were ranked on their over-expression upon *elt-2* depletion. (B) Percent of genes with a significant ELT-2 ChIP-seq peak in their promoter, split by intestine-enriched or not-intestine enriched status (black, ELT-2 promoter peak present; white, ELT-2 promoter peak absent). The total number of genes in each set is indicated below each bar. (C) Percent of genes that respond to *elt-2* depletion by down-regulation, up-regulation, or no response, split by ELT-2 binding status and intestine-enriched expression (green, ELT-2 activated; pink, ELT-2 repressed; white, ELT-2 independent). Percentages were measured for direct targets with ELT-2 promoter peak (top, ELT-2 present) and indirect targets without an ELT-2 promoter peak (bottom, ELT-2 absent). The total number of genes in each set is indicated below each bar.

**Figure 5.**
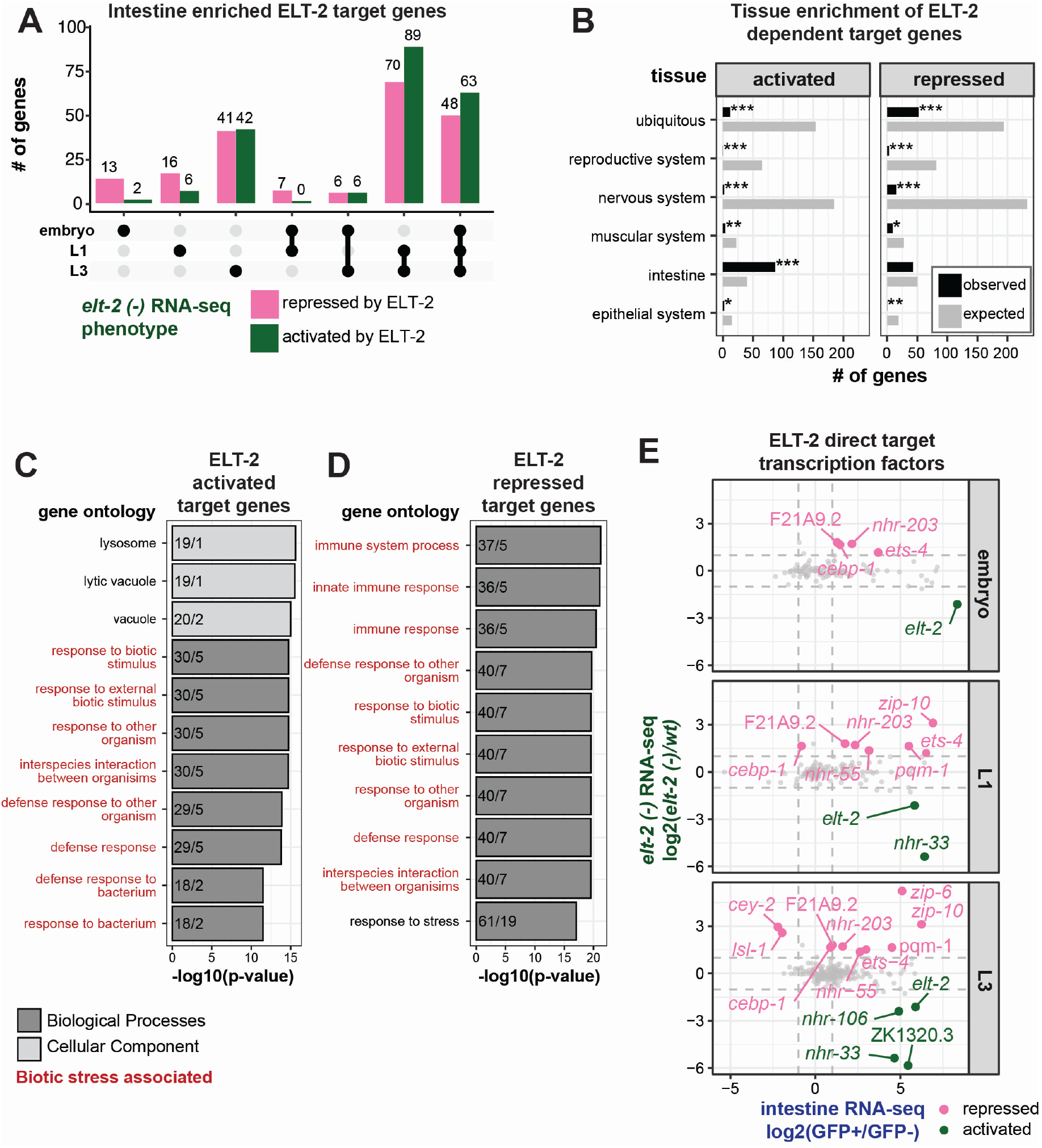
The direct targets of ELT-2 binding respond in diverse ways to *elt-2* loss. (A) The number of ELT-2-bound, intestine-enriched genes that respond to *elt-2* loss with up- or down-regulation are shown, split by developmental stage. The dots below the x-axis indicate gene set categories. Single dots are unique to one stage, while dots connected by lines indicate genes shared between stages. (B) Among genes that are purportedly activated or repressed by ETL-2, categories of genes associated with tissuespecific expression are not strongly over- or under-represented. The hypergeometric statistical test was used to test enrichment and depletion of gene set numbers for each category (black, observed number of genes; grey, expected number of genes; *P-value < 0.01, **P-value < 1×10^-5^, ***P-value < 1×10^-10^). (C-D) Gene Ontology (GO) analysis of direct ELT-2 activated (C) or repressed (D) gene sets across developmental stages. The top 10 GO terms for all three categories are displayed (BP, biological process; CC, cellular component). The x-axis corresponds to the −log10 transformed p-value, and the y-axis corresponds to the identified GO term. Numbers within each bar represent the number of “observed vs. expected” genes in the input set that correspond to the given GO term. Categorical labels are colored depending on whether they are unique or shared across the two categories. (E) The degree of ELT-2-dependence is plotted against intestine-enrichment for ELT-2 bound transcription factor targets. Dotted lines correspond the log2 fold change threshold of 1 used for differential expression significance testing. Genes with significant *elt-2 (-)* transcriptional response are indicated with genes names (pink, repressed; green, activated, Benjamini-Hochberg adjusted p-value < 0.01). Grey dots indicate transcription factors without significant response to *elt-2 (-)* (Benjamini-Hochberg adjusted p-value > 0.01)

**Figure 6.**
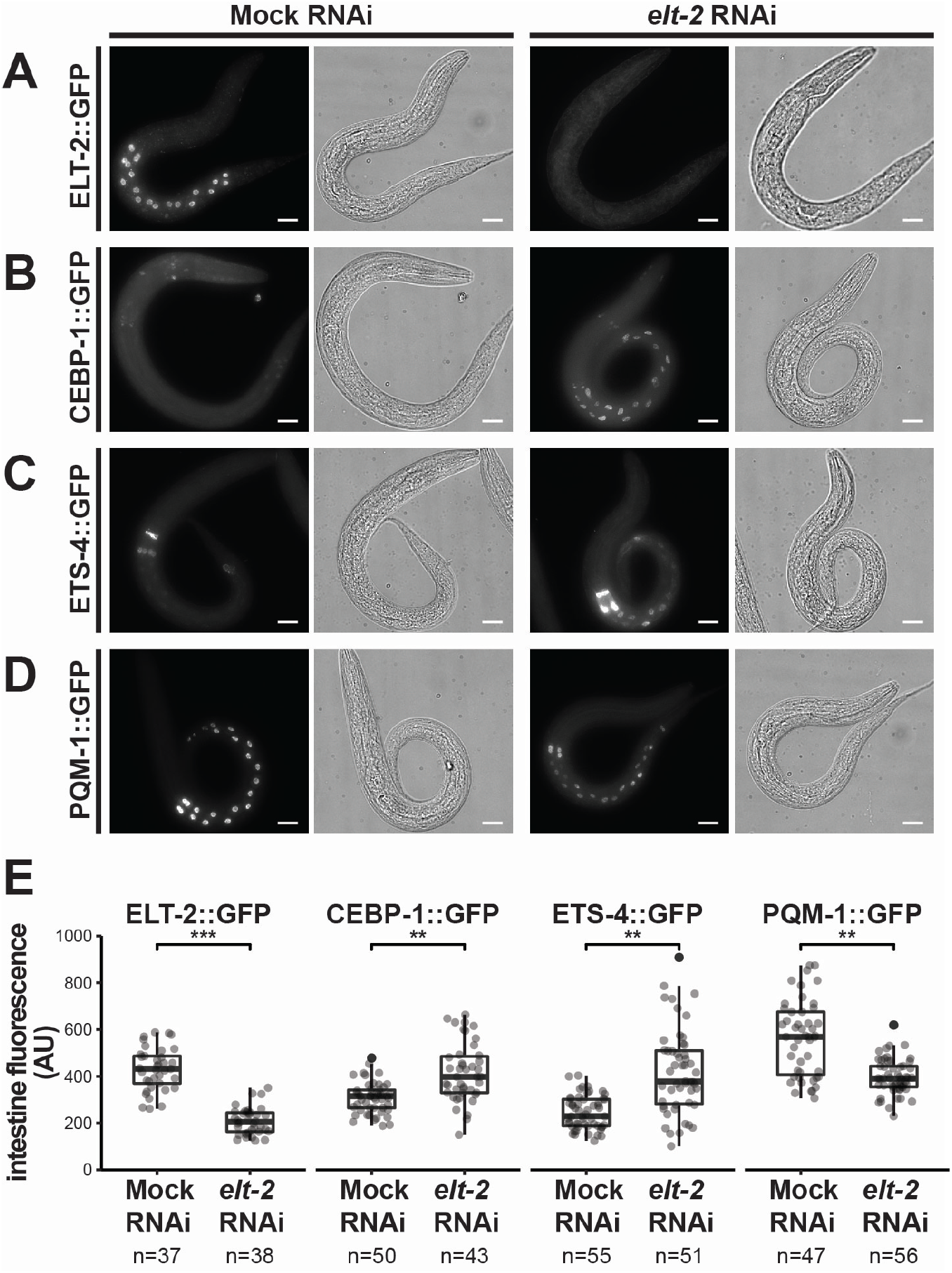
ELT-2 negatively regulates expression of transcription factors CEBP-1 and ETS-4 in the intestine. (A-D) *elt-2* RNAi depletion (right) compared to control (left) impacts transcription factor-GFP reporter strains. The strains imaged include: (A) ELT-2::GFP (*wgIs56 [elt-2::GFP]* allele, a control reporter to assess successful *elt-2* depletion), and query strains (B) CEBP-1::GFP (*wgIs563 [cebp-1::EGFP]* allele), (C) ETS-4::GFP (*wgIs509 [ets-4::EGFP]* allele), and (D) PQM-1::GFP (*wgIs201 [pqm-1::EGFP]* allele). To remove birefringent gut granule autofluorescence, animals were fixed and permeabilized through liquid nitrogen freeze-crack and subsequent washes with methanol, acetone, and PBST. Differential interference contrast microscopy (DIC) is also shown. Worms were imaged at 60x and multi-panel images were stitched as necessary. Scale bars = 10 microns. (E) Quantification of GFP signal in *C. elegans* intestines in GFP translation reporter strains (A-D). GFP measurements were collected from background subtracted maximum Z projections from three biological replicates. Intensity measurements were normalized for intestine area. Box and whisker plots represent the distribution of data points. Each point represents data measured from a single worm; filled points represent outliers (black). The total number of worms imaged are listed below the column label. Student’s t-test was calculated to measure statistical significance in GFP intestine fluorescence between mock and *elt-2* RNAi treatments (*P-value < 0.01, **P-value < 1×10^-5^, ***P-value < 1×10^-10^).

### RNAi feeding

To visualize the effect of ELT-2 depletion, we used *E. coli* feeding strains engineered to produce double-stranded RNA (dsRNA) cognate to the *elt-2* transcript. RNAi feeding strains were obtained from the Ahringer RNAi feeding library (Kamath and Ahringer 2003). Freshly starved worms were chunked to 150mm NGM/OP50 plates to grow for 48 hours. Synchronized embryos were isolated from mixed-stage populations through hypochlorite treatment (Stiernagle 2006). To visualize the effect of ELT-2 depletion on target gene expression (Figure 6), we grew synchronized embryos to the L3 stage by incubation at 20°C for 48 hours before RNAi exposure. L3 stage worms were transferred to NGM plates seeded with RNAi-inducing *E. coli* and incubated for an additional 48 hours until gravid. Synchronized L1s were collected by transferring RNAi treated gravid adults to 1 ml tubes of M9 and culturing for 24 hours. To image the response of *elt-2* promoter across developmental stages to ELT-2 depletion, we modified the above procedure by exposing worms to RNAi feeding strains 24 hours before the queried developmental stage (Figure 7, Figure S8). Negative control experiments utilized feeding strains containing L4440 empty vector. To confirm RNAi efficiency, we utilized POP-1 targeted RNAi as a positive control. RNAi experiments were considered successful when POP-1 RNAi resulted in 100% embryonic lethality. All RNAi plasmids were sequence verified before experimental use. Three biological replicates were performed for each GFP translation reporter strain.

**Figure 7.**
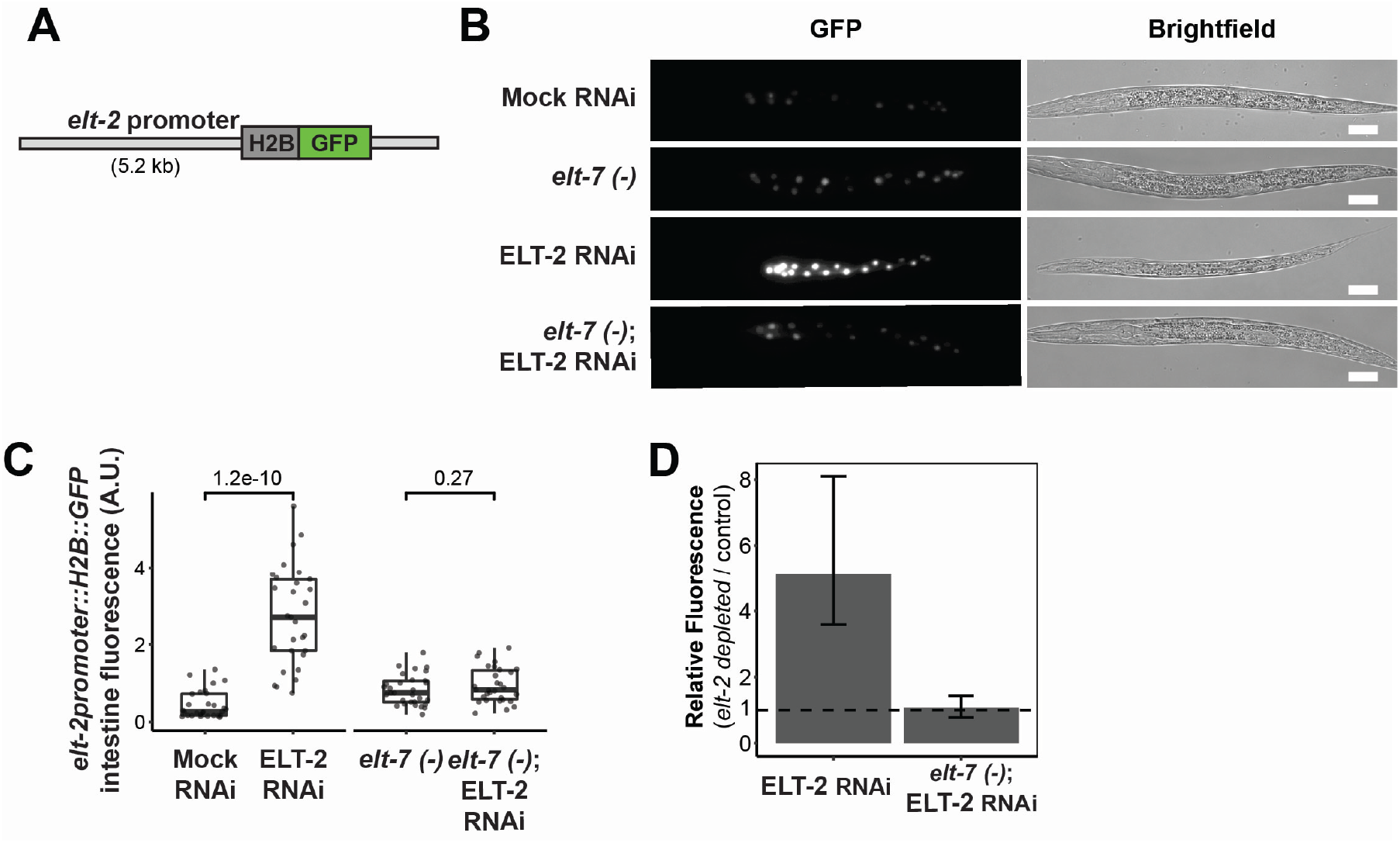
ELT-2 negatively regulates its own promoter. (A) Diagram of *elt-2* promoter reporter (*caIS71[elt-2p::H2B::GFP]* allele) used in this analysis. (B) Representative images of *elt-2* promoter activity in immobilized L1 stage animals. *elt-2* promoter reporter fluorescence was measured in *elt-7* wildtype or *elt-7 (tm840)* loss-of-function genetic background. Mock RNAi (L4440) or *elt-2* RNAi depletions were performed. Consistent imaging exposure was used between treatments and genetic backgrounds. Brightfield images are also shown. Animals were imaged at 40x. Scale bars = 20 microns. (C) Quantification of the *elt-2* promoter reporter’s fluorescence intensity. This analysis included intestine GFP signal quantification, background subtraction, and area normalization. Student’s t-test was used to determine statistical significance. Data represents 30 worms per treatment for three biological replicates. Box and whisker plots display the upper and lower quartiles while the middle line represents the median. Individual measurements are overlaid as points. (D) Comparison of *elt-2* promoter reporter activity for *elt-2* RNAi conditions with or without *elt-7* activity. Horizontal dotted line indicates relative fluorescence= 1. Error bars represent the t-test 95% confidence interval for the ratio of means between *elt-2* RNAi and control RNAi measurements.

### Fluorescence Microscopy

The response of GFP translation reporter gene fluorescence to ELT-2 depletion was visualized using fluorescence microscopy. RNAi-treated worms were imaged on a DeltaVision Elite inverted microscope equipped with an Olympus PLAN APO 60x (1.42 NA, PLAPON60XOSC2) objective, an Insight SSI 7-Color Solid State Light Engine, and SoftWorx software (Applied Precision) using 1 μm z-stacks. To remove birefringent gut granule autofluorescence, animals were fixed and permeabilized through liquid nitrogen freeze-crack and subsequent washes with methanol, acetone, and PBST.

For *elt-2* promoter experiments, images were collected using a Keyence BZ-X800 fluorescence microscope equipped with a Keyence PLAN APO 40x (0.95 NA, BZ-PA40) objective using 1 μm z-stacks. Worms were paralyzed with 25 mM sodium azide and mounted on microscope slides suspended in M9 and sealed with Valap. Imaging settings were kept constant between RNAi treatments and developmental stages per strain. Multi-panel images were processed and stitched, maximum Z projections were generated, and intestine fluorescence reporter was measured in ImageJ distribution Fiji (Preibisch et al. 2009; Schindelin et al. 2012).

Signal quantification was restricted to the intestine by utilizing a DIC reference image. Analysis and plots were generated using custom R scripts. Where ratios of reporter intensity are shown against controls with a p-value and confidence interval, a t-test of the ratio of means was performed using the ttestratio function in R package mratios (v 1.4.2), with the null hypothesis of a ratio of 1. Confidence intervals are 95% standard error around the ratio of means (Djira et al. 2020).

### Single molecule inexpensive fluorescence *in situ* hybridization (smiFISH)

smiFISH was used to confirm intestine specificity of bioinformatically identified intestine genes. Transcripts were probed in the embryo stage using smiFISH as previously described (Tsanov et al. 2016; Parker et al. 2021). Probes were designed with FLAP X extensions for *acy-4, bre-2*, and C02D5.4 and FLAPY for *ges-1* (File S8). For each gene 12 primary probes were designed with the R script Oligostan (https://bitbucket.org/muellerflorian/fish_quant/src/master/Oligostan/, commit d87b4b2). Secondary FLAP probes were ordered from Stellaris LGC with dual 5’ and 3’ fluorophore labeling. FLAPY probes were labeled with Cal Fluor 610 and FLAP X probes were labeled with Quasar 670 (Biosearch Technologies, BNS-5082 and FC-1065, respectively). All samples were imaged on a DeltaVision Elite inverted microscope described above. Images were processed in the ImageJ distribution Fiji (Schindelin et al. 2012).

## Results

### *C. elegans* intestine transcriptional profiling by FACS isolation in embryonic and post-embryonic stages

We assessed the intestine-specific transcriptome across three stages: embryo, L1, and L3. Previous works had characterized intestine-specific transcriptomes but either at a single stage per study or using a mix of stages *en masse*. Those works employed single-cell dissection of embryonic cells, RNA immunoprecipitation techniques, and intestine purification through Fluorescence Activated Cell Sorting (FACS), primarily in the embryo, L1 or L2 stages (Spencer et al. 2011; Haenni et al. 2012; Blazie et al. 2015; Hashimshony et al. 2015; Tintori et al. 2016; Kaletsky et al. 2018). Technical differences across the studies prevented their integration across datasets. Therefore, a comprehensive view of intestine gene expression over developmental time had been missing.

Here, we have taken a FACS-based approach to isolate intestine cells for RNA-seq analysis (Figure 1A). Intestine cells were labeled with the transgenic intestinal reporter *elt-2p::H2B::GFP* (Figure 1B). Worm dissociation protocols and FACS gating strategies were optimized for each of the three stages (see Materials and methods, Figure S1). A flow cytometry profile of dissociated cells detected 2% of cells as GFP-labeled, which corresponds to the expected percent of intestine cells in the worm (20/ 1000 = 2%). GFP-labeled intestine cells were then isolated by FACS with a purity of 80-90% (Figure 1C) and verified visually for GFP fluorescence (Figure 1D).

RNA-seq libraries were generated on intestine cell (GFP+) and non-intestine cell (GFP-) populations (File S1-S3). To determine the similarities and differences between samples on a genome-wide scale, we performed sample-to-sample correlation analysis (Figure S2). This analysis identified that samples clustered by stage and GFP fluorescence. We were concerned that the process of cell dissociation itself could introduce transcriptional changes due to cell stress or handling. To evaluate the extent of these changes, we determined the transcriptional differences between whole worms and freshly dissociated but unsorted cells (Figure S3A). We found that cell dissociation imposed only minimal changes in the embryo stages and some moderate differences in the larval stages. Genes that significantly changed due to cell dissociation were filtered from downstream analyses.

To determine whether our intestinal RNA-seq samples were indeed enriched for transcripts of known intestine-expressed genes, we compared the GFP+ to GFP- transcript abundance for genes specific to the intestine (*act-5, elt-2, ges-1, ifb-2*), germline (*glh-1, meg-1, pgl-1*), neuron (*rab-3, rgef-1, rgs-1*), muscle (*fhod-1, hlh-1, myo-3*), and hypodermis tissues (*bli-1, lon-3*) (Figure 1E). In all assayed developmental stages, only intestine-specific genes scored a significant fold-change score for enrichment in the isolated GFP+ intestine cells, using DESeq2. Of the 11 non-intestine genes investigated, none were enriched in the intestine. This result confirms that our FACS-isolated transcriptomes indeed represent the intestine.

### Differential RNA-seq analysis detects intestine-enriched transcripts

To identify intestine-expressed transcripts in each of the embryo, L1, and L3 stages, differential expression analysis was performed using DESeq2 (Figure 2A - C). In each of the three stages of development, roughly 1,500 intestine-enriched genes were identified. When we compared across developmental stages (Figure 2D), we found a large set of these genes were intestine-enriched in all three stages (n = 848), indicating a core set of intestinal genes. Genes specific to the embryonic stage comprised the next largest gene set (n = 644), suggesting that gene expression in the embryonic intestine is distinct from larval expression.

Gene Ontology (GO) analysis identified categories of genes that were shared in all stages of intestine-enriched genes such as “innate immune response”, “peroxisome”, “membrane raft”, “carbohydrate binding”, and “glucuronosyltransferase activity”. These categories are consistent with known intestinal biological functions that occur in the intestine (Figure 2E-F). GO terms associated specifically with the embryo stage included DNA-binding transcription factor activity, indicating an overrepresentation of transcriptional regulators. GO terms associated specifically with the L1 stage include “proton-transporting V-type ATPase complex”, and terms associated specifically with the L3 stage include “defense response to Gram-negative bacterium”, suggesting the importance of defense at this stage. Again, these findings reinforce the result that some gene expression tasks are common across all developmental stages whereas others are unique to a given stage.

To validate our intestine-enriched RNA-seq dataset, we utilized smiFISH (single molecule inexpensive Fluorescence In Situ Hybridization) (Tsanov et al. 2016; Parker et al. 2021). We were interested in determining if our dataset could identify previously uncharacterized intestine-enriched transcripts by microscopic visualization. The transcripts *acy-4, bre-2*, and C02D5.4 were detected as intestine expressed across all stages in our study but were not annotated as intestine-enriched in WormBase (Figure 3A). We assessed their mRNA abundance and localization in embryos while co-staining for the intestine marker transcript *ges-1* (Kennedy et al. 1993). *acy-4* and *bre-2* mRNA were exclusive to intestine tissue (Figure 3B, C). C02D5.4 mRNA localized to the intestine and epithelial cells yet still appeared intestine-enriched compared to the embryo as a whole (Figure 3D). This illustrates that our intestine-specific transcriptome profiling was sufficient to identify new intestine-enriched genes.

### Integration of genome-wide datasets

Previous reports have proposed that ELT-2 participates in the transcription of every gene expressed in differentiating and mature intestines (McGhee et al. 2007, 2009). This assertion was based on the high prevalence of ELT-2 binding site sequences (TGATAA) in intestine promoters. However, ChIP-seq data across many fields illustrates that only a fraction of possible binding sites are typically occupied by TFs, usually due to either chromatin context, combinatorial binding with other TFs, or other causes (Ucar et al. 2009). To understand the scope of ELT-2’s influence in the developing intestine GRN, we integrated 1) previously published ELT-2 binding maps (ELT-2 ChIP-seq; File S4) (Kudron et al. 2017), 2) our intestine-specific RNA-seq profiles (intestine RNA-seq, GFP+/GFP-), and 3) whole transcriptome responses to elt-2 deletion (RNA-seq) (Dineen et al. 2018) (Figure 4). We divided the data into subsets based on ELT-2 ChIP-seq occupancy (ELT-2 bound) and RNA-seq expression status within the intestine (intestine-enriched) (Figure 4, File S5). “ELT-2 bound” genes contained ELT-2 ChIP-seq peak within 1 kb upstream and 200 bp downstream of their transcription start site. Genes were “intestine-enriched” if they scored an intestine vs. non-intestine fold-change greater than 1 and p-value < 0.01 (see Methods). We plotted all three datasets as aligned heatmaps, filtering for all protein-coding genes in the genome. Finally, we used the transcriptional response to *elt-2* loss (plotted as fold-change ratio and sorted within categories based on this response) to order genes within each category. Genes dependent on ELT-2 for their expression (activated by ELT-2; i.e: they are under-expressed upon *elt-2* depletion) are shown in green, and genes that are conversely repressed by ELT-2 (they become over-expressed upon *elt-2* depletion) are shown in pink.

It should be noted that the study assessing transcriptional responses to *elt-2* depletion was collected by COPAS sorting whole L1 worms that had either lost or retained an unstable ELT-2::GFP rescue array (Dineen et al. 2018). Therefore, that dataset does not include embryo- or L3-specific responses. Also, because the data was collected in whole worms, it may capture responses occurring outside of the intestine such as indirect responses in other tissues. However, this dataset still provides useful information regarding the role of ELT-2 in the intestine GRN.

### Quantification of ELT-2 target genes in the intestine GRN

Previous work hypothesized that ELT-2 directly regulates the expression of most intestine genes owing to a prevalence of the ELT-2 binding site within intestinal promoters (McGhee et al. 2009). If ELT-2 occupancy is a major predictor of intestine-enriched expression, we would expect to see significant overlap between “ELT-2 bound” and “intestine-enriched” categories (Figure 4). This would suggest that ELT-2 occupancy is correlated with intestine enrichment and could be predictive of it. In contrast, if ELT-2 occupancy is not a major predictor of intestine-enriched expression, we would expect that genes with “ELT-2 bound” promoters would fail to yield “intestine-enriched” annotations. To test this, we tabulated the ELT-2 bound and intestine-enriched categories (Figure 4B). In all stages, ELT-2 bound genes were more likely to be intestine-enriched than genes not bound by ELT-2 (chi-square p-value < 1E-100), suggesting that ELT-2 occupancy does assist in predicting intestine-enriched expression. In the embryo stage, however, only 31% (572/1745 genes) of intestine-enriched genes were bound by ELT-2, indicating that ELT-2 occupancy alone is insufficient to predict a gene’s intestine-enriched status. This percentage increased in L1 (55%, 872/1485 genes) and L3 (75%, 1130/1445 genes) stages (Figure 4B), illustrating that ELT-2 may be responsible for the regulation of a larger fraction of larval stage intestinal transcriptomes than embryonic ones. Overall, these results suggest intestine enrichment implies ELT-2 binding, but that not all intestine-enriched genes are bound by ELT-2, particularly in the embryo stage.

### Intestine-enriched gene expression as a function of ELT-2 ChIP signal

Our findings illustrate that a subset of ELT-2 binding is associated with intestine-enriched gene expression. For this subset, our next question was whether the degree of ELT-2 occupancy could predict the degree of transcriptional output. A simple mechanistic model would predict that transcription factor binding quantitatively influences transcriptional output such that genes with high ELT-2 occupancy would be highly intestine expressed. In contrast, an alternative model could posit that ELT-2 occupancy is independent from the degree of intestine expression.

To distinguish between these scenarios, we first plotted intestine RNA-seq counts (regularized and log-transformed) for ELT-2 bound versus not bound genes (Figure S4, File S7). For all developmental stages, ELT-2 bound genes, as a population, had significantly higher intestine RNA-seq counts than not bound genes indicating that ELT-2 occupancy is associated with higher overall expression. Still, genes not bound by ELT-2 achieved a similar range of expression in the intestine, indicating that other factors independent of ELT-2 binding can also promote high intestinal gene expression.

We next assessed whether the degree of ELT-2 occupancy was correlated with the degree of gene expression by plotting RNA-seq counts over the ELT-2 ChIP-signal (Figure S5A). The R2 values ranged from 0.04 – 0.14, indicating low predictive value and suggesting against a scenario where the degree of ELT-2 occupancy informs gene expression level. Stratifying based on ELT-2 regulatory status (Figure S5B) failed to improve the correlation coefficients, again indicating a low predictive value.

Though we failed to link ELT-2 occupancy to expression level, we did notice that ELT-2 occupancy is dynamic over developmental time. Plotting the averaged ELT-2 ChIP-seq signal across meta-promoter regions for different expression categories (Figure S6), we observed that ELT-2 occupancy was greater across the promoters of intestine-enriched genes than non-intestine genes in L1 and L3 stages but not in embryos. This corroborates our previous findings that ELT-2’s relative contribution to the intestinal transcriptome increases over developmental time (Figure 4B).

Altogether, the degree of ELT-2 occupancy does not instruct the degree of transcriptional output. Furthermore, the degree of ELT-2 occupancy is dynamic over developmental time, increasing in both signal intensity and breadth in L1 and L3 intestinally expressed genes. That is, ELT-2 influences a greater proportion of L1 and L3 intestinal gene expression. In contrast, intestine expression in embryos may be achieved through alternative transcription factors, as yet unidentified.

### ELT-2 regulates target genes through activation and repression

Previous reports have suggested that ELT-2 functions exclusively as an activator of the intestine gene regulatory network (McGhee et al. 2007; Wiesenfahrt et al. 2015; Lancaster and McGhee 2020). To evaluate if ELT-2 serves to primarily activate target gene transcription, we integrated RNA-seq data measuring differentially expressed genes in *elt-2 (-)* versus wildtype conditions (Figure 4C). Of the genes that are both ELT-2 bound and intestine-enriched, half did not depend on ELT-2 alone to maintain their proper transcriptional level (embryo 61.6%, L1 44%, L3 46%). Of those that did experience gene expression changes upon *elt-2* depletion, an equal number were positively regulated by ELT-2 (ELT-2 activated: embryo 18.6%, L1 29.3%, L3 29.0%) as were negatively regulated (ELT-2 repressed: embryo 19.6%, L1 26.0%, L3 24%).

### ELT-2 represses defense response genes

Since ELT-2 is intestine expressed throughout development, we wondered how ELT-2’s role as an activator or a repressor of downstream target genes varied over time. We observed 13 ELT-2 repressed genes and 2 ELT-2 activated genes distinct to the embryonic stage (Figure 5A). Conversely, we observed 48 ELT-2 repressed genes and 63 ELT-2 activated genes shared between all assayed stages. This suggests that there does not appear to be a heavy bias in the developmental stage for either ELT-2 repression or activation activity in the intestine GRN. Furthermore, we identified 70 ELT-2 repressed genes and 89 ELT-2 activated genes shared between L1 and L3 stages. Together, these results suggest that ELT-2 has a developmentally dynamic role in the intestine GRN, but the molecular factors that dictate stage-specific ELT-2 regulatory dynamics are unknown.

Previous reports of human GATA factors have demonstrated that they promote one fate while repressing an alternative fate (Tremblay et al. 2018). We were interested in determining if ELT-2 functions this way. To evaluate this, we generated nonoverlapping sets of anatomy ontology terms for major organ systems, including reproductive system, nervous system, muscular system, intestine, and epithelial system, and a separate set of ubiquitously expressed genes (File S6) (Rechtsteiner et al. 2010). We used hypergeometric statistics to evaluate the enrichment or depletion of terms for all ELT-2 target genes from all stages stratified based on ELT-2 activation or repression (Figure 5B). ELT-2 activated target genes were significantly enriched in genes associated with intestine and significantly depleted in genes associated with both non-intestine and ubiquitous genes. ELT-2 repressed target genes were not over-represented for any of the categories we assayed suggesting that ELT-2 does not play a clear role in repressing an alternative tissue type. That is, ELT-2 activated genes are strongly associated with intestine genes, but ELT-2 repressed genes are not associated with any one tissue. It is possible that ELT-2 repressed genes are intestine-expressed genes that are normally repressed by ELT-2 and only induced under certain conditions.

We were curious whether the genes that ELT-2 represses shared similar functions suggesting that repression by ELT-2 could prevent specific biological characteristics. To evaluate this, we performed GO analysis on ELT-2 activated or repressed direct target genes identified from all three developmental stages assayed (Figure 5C, D). ELT-2 activated target genes were associated with defense response to bacterium and related lysosome terms. ELT-2 repressed target genes were associated with response to stress and related innate immune response terms. Both ELT-2 activated and repressed target genes were associated with several related defense response terms and interspecies interaction between organisms. Overall, these results suggest that both ELT-2 activation and repression activity is central in regulating responses to biotic stimulus. Our results suggest that ELT-2 turns on lysosome and a set of immune-related genes, whereas ELT-2 repression is central to regulating innate immunity genes. These findings are consistent with previous reports which have demonstrated ELT-2 regulation of lysosome and immunity pathways (Block et al. 2015; Keith et al. 2016).

Overall, our results suggest that ELT-2 is not alone in driving intestine-enriched gene expression, particularly in embryonic stages. Other TFs contribute independently or downstream of ELT-2 action. For those TFs that are downstream of ELT-2, we were curious which were likely to be ELT-2 activated or ELT-2 repressed. To determine this, we subset ELT-2 regulated target genes for known and predicted TFs (Figure 5E) (Bass et al. 2016). We identified a set of transcription factors activated by ELT-2 (*nhr-33, nhr-106*, and ZK1320.3) and a set repressed by ELT-2 (*cebp-1, ets-4, nhr-203*, F21A9.2, *nhr-55, zip-10, pqm-1, cey-1*, and *lsl-1). elt-2* was detected as an activated target due to the nature of the *elt-2* deletion allele used in this study. The number of ELT-2 regulated target transcription factors increased over developmental time. ELT-2 regulation of these transcription factors may explain the observed ELT-2 transcriptional dependence for genes without evidence of ELT-2 promoter binding (Figure 4C). These results further emphasize that ELT-2 regulates the intestine gene regulatory network through both activation and repression of target genes. This information will be helpful to set the stage for future studies aimed at expanding the intestinal GRN to encompass more numerous TFs.

Genes that were not bound by ELT-2, still respond to *elt-2* depletion albeit at a lower rate (Figure 4C). In all cases, we identified a statistically significant relationship between intestine enrichment and the distribution of ELT-2 regulated gene states (chi-square p-value < 1E-35). These results suggest that the bulk of direct ELT-2 targets are not dependent on ELT-2 for their transcriptional activity. It is possible that other factors or contexts may be required for ELT-2 transcriptional regulation. Additionally, these results demonstrate that ELT-2 functions not only as an activator but also represses distinct sets of target genes.

### ELT-2 negatively regulates the expression of transcription factors CEBP-1 and ETS-4 in the intestine

Our analysis suggests that ELT-2 regulates the intestinal GRN through both activation and repression. Previous work has focused on ELT-2 as a transcriptional activator of its direct targets and transcriptional evidence supports the idea that ELT-2 may also act as a transcriptional repressor of certain direct targets, but with no definitive direct evidence (Dineen et al. 2018). It is possible that ELT-2 may either downregulate gene expression through direct repression or indirectly through complex feedback loops. Additionally, it is possible that some genes that become over-expressed upon ELT-2 depletion are the result of over-expression in other tissues indicating an indirect and non-intestinal response. To better assess ELT-2’s negative regulatory behavior and to test whether this regulatory behavior is centered within the intestine, we selected three genes that are the direct targets of ELT-2 binding and that were up-regulated upon *elt-2* deletion: *cebp-1* (CCAAT/enhancer-binding protein), *ets-4* (ETS class transcription factor) and *pqm-1* (ParaQuat Methylviologen responsive) (Figure S7A-C). We assessed whether GFP translation fusion constructs for these TFs recapitulated the overexpression phenotype observed upon *elt-2* deletion in RNA-seq data. We simultaneously evaluated whether over-expression was intestine-specific. We depleted ELT-2 by RNA interference (RNAi), and ELT-2 knockdown was validated by observing a significant 2-fold reduction in ELT-2::GFP signal in intestinal nuclei compared to control worms at matched developmental stages (p-value = 1.2E-18) (Figure 6A, E).

*cebp-1* encodes a bZIP (basic leucine zipper) class transcription factor with annotated expression in the intestine, hypodermis, pharynx, muscle, and neuronal cells. *cebp-1* is essential for neuronal axon regeneration, stress response, and intestinal immune response (Yan et al. 2009; McEwan et al. 2016; Yang et al. 2017; Wu et al. 2021). In native ELT-2 conditions, CEBP-1::GFP fluorescence was observed in pharyngeal nuclei. Depletion of ELT-2 led to a significant 1.4-fold increase of CEBP-1::GFP reporter activity in intestine nuclei (p-value = 3.3E-6) (Figure 6B, E) further confirming that the over-expression response is specific to the intestine.

*ets-4* encodes an ETS (E26 transformation-specific) class transcription factor with a winged helix-turn-helix DNA binding motif (Sharrocks 2001). *ets-4* has annotated expression in the intestine, germline, adult hypodermis, and seam cells and functions in lifespan regulation and axon regeneration (Hart et al. 2000; Thyagarajan et al. 2010; Li et al. 2015; Kaymak et al. 2016; Sakai et al. 2019). In native ELT-2 conditions, we observed ETS-4::GFP reporter activity localized to the nuclei of the pharyngeal-intestinal valve (vpi) cells, anterior intestine nuclei, and rectal gland cells (Figure 6C). Upon ELT-2 depletion by RNAi, we observed a significant 1.7-fold increase of ETS-4::GFP reporter activity in intestine nuclei compared to mock RNAi controls (p-value = 1.3E-8) (Figure 6C, E). Similar to CEBP-1, ETS-4 therefore responds to ELT-2 loss with intestine-specific ETS-4 over-expression.

*pqm-1* is a C2H2-type zinc finger transcription factor that is primarily expressed in the intestine and neuronal cells (Deplancke et al. 2006; MacNeil et al. 2015; Bass et al. 2016; O’Brien et al. 2018). *pqm-1* functions in several processes including stress response, defense response, lipid metabolism, and lifespan regulation (Shapira et al. 2006; Tepper et al. 2013; O’Brien et al. 2018; Dowen 2019; Heimbucher et al. 2020). We observed PQM-1::GFP reporter signal localized to intestine nuclei (Figure 6D). In contrast to ETS-4::GFP and CEBP-1::GFP, ELT-2 depletion had an opposite effect on PQM-1::GFP reporter activity than expected. Intestine localized PQM-1::GFP reporter activity was significantly reduced 1.5-fold when ELT-2 was depleted by RNAi (p-value = 1.4E-8) (Figure 6D, E).

The increase in intestine-localized GFP for CEBP-1::GFP and ETS-4::GFP upon ELT-2 depletion is consistent with the model that ELT-2 functions to directly repress these direct target genes. We selected these transcripts for hypothesis validation because we detected an ELT-2 peak in their promoters (Figure 5E, Figure S7A, B). Our reporter assay results illustrate that both CEBP-1 and ETS-4 depend on direct ELT-2 regulation to remain lowly expressed in the intestine. It remains possible that in addition to ELT-2’s direct action at these promoters that indirect negative feedback loops are also involved. In contrast, the evidence that PQM-1::GFP protein abundance decreases upon ELT-2 depletion is incongruous with the evidence that *pqm-1* transcript abundance increases upon ELT-2 depletion (Figure 5E, Figure S7C). This result does not support our initial hypothesis for the regulatory connection between ELT-2 and PQM-1 but does demonstrate that PQM-1 is dependent on ELT-2 genetic control. This suggests that PQM-1 is not controlled by a simple ELT-2 transcriptional repression model and that PQM-1 may be under more complex genetic control than anticipated, possibly with transcriptional and translational regulatory components. Additionally, this result emphasizes previous findings that mRNA transcript abundance is often insufficient to predict protein abundance (Gygi et al. 1999; Bauernfeind and Babbitt 2017).

That *cebp-1* and *ets-4* are upregulated in the absence of *elt-2* supports our hypothesis that *elt-2* plays a repressive role in their transcriptional regulation. The combined evidence that ELT-2 directly binds their promoters and is required for their downregulation suggests that ELT-2 represses these target genes. However, the molecular mechanisms regulating this process require further experimentation to assess whether this is due to recruitment of transcriptionally repressive machinery or due to a complex negative regulatory feedback loop.

### ELT-2 negatively regulates its own promoter

ELT-2 binds its own gene promoter (Fukushige et al. 1999; Wiesenfahrt et al. 2015). The reported purpose of this regulation is to sustain ELT-2 expression throughout the worm’s lifetime through autoactivation until it declines in old age (Mann et al. 2016). Previous transcriptomics studies were unable to discern how *elt-2* promoter output responded to *elt-2* depletion, as the studies were conducted in *elt-2* deletion backgrounds (Dineen et al. 2018). Having detected ELT-2 repressive activity, we thought it pertinent to reevaluate how ELT-2 impacts its own promoter output. By using an *elt-2* promoter reporter transgene driving histone H2B fused to GFP, we could report *elt-2* promoter activity divorced from the production of ELT-2 protein (Figure 8A). If ELT-2 positively regulates its own promoter’s expression, we expect ELT-2 depletion to reduce *elt-2* promoter activity. Surprisingly, depletion of ELT-2 protein in the L1 stage resulted in a 5-fold over-expression of *elt-2* promoter-driven GFP compared to mock RNAi controls, suggesting that ELT-2 negatively regulates *elt-2* promoter activity (Figure 7B-D). We observed this negative autoregulation in L1, L3, and adult stages suggesting that this activity is not restricted to one developmental stage (Figure S8).

**Figure 8.**
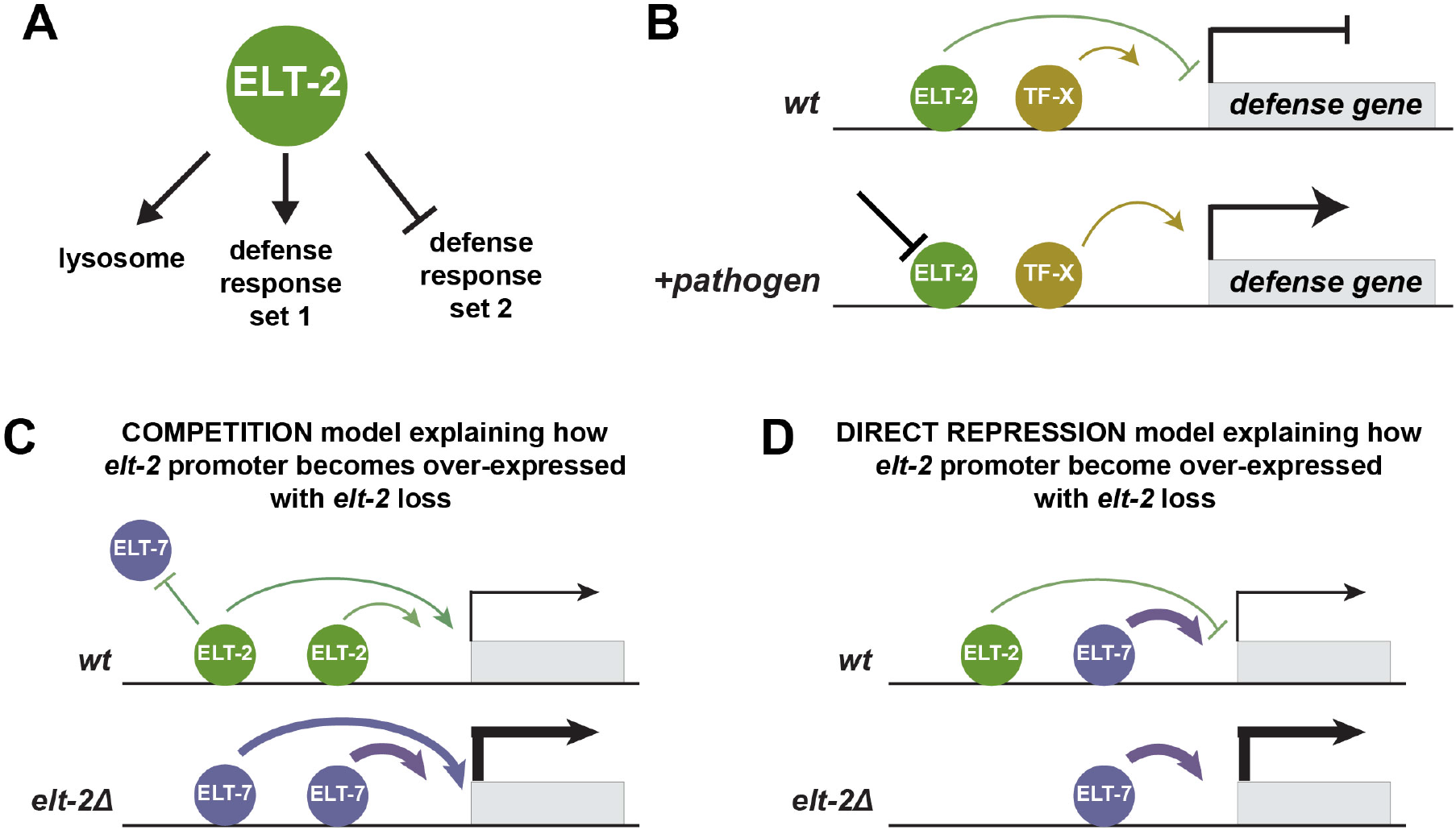
Model summarizing ELT-2 regulatory activity. (A) ELT-2 promotes lysosome and one set of defense response genes while simultaneously repressing a distinct set of defense response genes. (B) We hypothesize that ELT-2 repression of defense-response genes are inducible in situations such as pathogen exposure where relief of ELT-2 repression is required for defense response gene expression. (C) ELT-2 performs negative autoregulation with apparent ELT-7 overcompensation. We hypothesize two scenarios explaining this result. Either ELT-2 and ELT-7 are directly competing for binding to the *elt-2* promoter and ELT-7 serves as a stronger activator than ELT-2, or (D) ELT-2 is directly repressing the *elt-2* promoter whereas ELT-7 activates it.

Roughly half of genes that are over-expressed in the absence of *elt-2* do so in manner dependent on *elt-7*, an *elt-2* homolog. These are members of the previously identified *“elt-7* over-compensated” gene set (Dineen et al. 2018). Previous studies have been unable to determine how ELT-2 responds to *elt-2* and/or *elt-7* deletion as the *elt-2 (ca15)* allele previously used to perform transcriptomics was a complete coding sequence deletion. The use of this allele obscured any transcriptional response of *elt-2* to either *elt-2* or *elt-7* mutations. To determine if *elt-2* is also an “*elt-7* over-compensated” gene, we measured *elt-2* promoter activity in an *elt-7* genetic deletion background. We observed that the *elt-2* promoter is no longer over-expressed when both ELT-2 and ELT-7 are depleted (Figure 7B-D), thereby demonstrating that ELT-2 is also “*elt-7* over-compensated”. In other words, in the absence of ELT-2, the *elt-2* promoter becomes over-activated in an ELT-7-dependent fashion.

We were interested to determine if our results demonstrating ELT-2 negative autoregulation were specific to the reporter construct or if these phenomena could be observed using a complementary method. To do this, we re-analyzed our RNA-seq data and visualized the *elt-2* genomic locus for RNA-seq reads in *elt-2 (-)* mutants with or without *elt-7* (Figure S9A). We measured increased reads aligning to the *elt-2* promoter in *elt-2 (-)* mutants compared to wildtype (Figure S9B). Additionally, the increase in *elt-2* promoter-localized RNA-seq reads were reduced in *elt-2(-);elt-7(-)* double mutants. These results serve to support the biological significance of our findings in a reporter construct context.

Together, these results suggest that ELT-2 protein negatively regulates its own promoter in an ELT-7 dependent manner. We suggest that ELT-7 is responsible for over-activating the *elt-2* promoter when relieved of ELT-2 repression. Previous transcriptomics work has shown that *elt-7* is not over-expressed upon loss of *elt-2* at the mRNA level (Dineen et al. 2018), suggesting that the higher activity of ELT-7 on the *elt-2* promoter is post-transcriptional, potentially occurring through an increased ability of ELT-7 to stimulate transcriptional output. Intriguingly, the *elt-2* promoter is still expressed at wildtype levels when both ELT-2 and ELT-7 are absent, implying that an additional factor may contribute to *elt-2* transcription (Figure 7C, D). These findings demonstrate that the wild-type level of *elt-2* mRNA production is affected by both positive and negative inputs. This example emphasizes combinatorial control by both ELT-2 and ELT-7 in regulating the intestine gene regulatory network and illustrates that an additional unknown factor (or factors) participate(s) in the intestinal GRN.

## Discussion

ELT-2 is the major transcription factor that marks intestinal tissue. ELT-2 production commences in embryos where its role is well studied, yet it remains present during larval and adult stages where its impact is more nebulous. *C. elegans* larval and adult-stage intestines perform digestion and metabolism characteristic of most animal intestines. However, they also perform additional functions that in other animals are overseen by separate organs such as yolk production, insulin signaling, regulation of developmental aging, immune response, stress response, and detoxification (Kimble and Sharrock 1983; An and Blackwell 2003; Libina et al. 2003; Martinez-Finley and Aschner 2011; Ludewig and Schroeder 2013; Head and Aballay 2014; Block et al. 2015; Keith et al. 2016; Mann et al. 2016; Chun et al. 2017; Lee and Mylonakis 2017; Zárate-Potes et al. 2020).

Though ELT-2 is required for optimal performance of many of these sub-functions, the intestinal GRN has yet to be mapped to derive how ELT-2 influences them. In this study, we used a systems biology approach to place ELT-2 within the larger GRN and our findings countered previous assumptions that 1) ELT-2 binding is associated with the expression of all intestine genes, 2) ELT-2 functions solely as a transcriptional activator, and 3) ELT-2 performs positive autoregulation.

Previous work suggested that ELT-2 is predominantly responsible for intestine gene activation due to a high prevalence of TGATAA sequences in intestine gene promoters and based on the assumption that ELT-2 behaves as an activator (Fukushige et al. 1998, 2003, 2005; Kalb et al. 2002; Oskouian et al. 2005; McGhee et al. 2007, 2009; Sommermann et al. 2010). Here, we combined ELT-2 binding maps (ChIP-seq), transcriptome response to *elt-2 (-)* (RNA-seq), and our newly generated intestine transcriptome data to evaluate this hypothesis. We identified that 31% of embryo stage intestine-enriched genes had observable ELT-2 binding in their promoters and that the percent of intestine-enriched genes bound by ELT-2 increased over developmental time to 75% in the L3 stage. This suggests that not all intestine-enriched genes are regulated by ELT-2 but that the intestinal transcriptome becomes more reliant on ELT-2 in later developmental stages. Further analysis of the intestine transcriptome data generated in this study may be of benefit for identifying additional intestine GRN components, particularly in the embryonic stages.

ELT-2 has largely been assumed to function predominantly as a transcriptional activator (McGhee et al. 2007, 2009). However, roughly half of the genes directly bound by ELT-2 are unaffected by *elt-2* loss. It is possible that these ELT-2 target genes may be redundantly regulated by other transcription factors, such as ELT-7. Surprisingly, of the ELT-2 direct target genes that do change in their transcript abundance upon *elt-2* loss, an equal proportion become up-regulated as down-regulated. This result is contrary to the prevailing hypothesis that ELT-2 functions solely as an activator (McGhee et al. 2007, 2009). As expected, GO analysis demonstrated that ELT-2 activated target genes are associated with lysosome and defense response processes (Figure 8A). We hypothesized that ELT-2 repression could serve to prevent alternative tissue type characteristics (like those associated with neurons or gonads). However, we found this was not supported by GO evidence. Categories associated with other tissue or cell types were not over-represented among the ELT-2 repressed targets. Instead, ELT-2 repressed target genes were strongly associated with innate immunity and defense response genes. We hypothesize that ELT-2 establishes a baseline expression of defense genes while also repressing other defense response genes, keeping them in balance until pathogen exposure (Figure 8B). Molecular details regulating this process will need to be experimentally determined (Shapira et al. 2006).

Of the ELT-2 repressed genes, we verified that two transcription factors, *cebp-1* and *ets-4*, were negatively regulated by ELT-2 specifically within the intestine. This refuted the possibility that their over-expression (upon *elt-2* loss) was occurring elsewhere in the worm as a downstream, indirect response. By placing their over-expression phenotypes squarely within the intestinal cells where ELT-2 is present, this result adds greater support to the idea that ELT-2 is intimately associated at a mechanistic level with their repression. *cebp-1* and *ets-4* are transcription factors that function in pathogen response with additional roles in neuronal axon regeneration (Yan et al. 2009; Thyagarajan et al. 2010; Li et al. 2015; McEwan et al. 2016; Malinow et al. 2019; Sakai et al. 2019; Wu et al. 2021). This result is consistent with our hypothesis that ELT-2 repression likely serves to keep defense response genes off until a stimulus-triggered relief of ELT-2 repression, consistent with previous reports (Block and Shapira 2015; Block et al. 2015). It is possible that negative regulation by ELT-2 could occur through negative feedback loops within the intestinal GRN or by recruiting repressive machinery (Figure 8A, B). Further studies will be required to differentiate between these models.

Having identified that ELT-2 functions as both an activator and repressor, we were interested in reevaluating the prevailing hypothesis that ELT-2 performs positive autoregulation. Previous work established that ELT-2 binds its own promoter through microscopy, ChIP-seq, and *in vitro* binding assays (Hawkins and McGhee 1995; Wiesenfahrt et al. 2015). The role of this binding was posited to maintain ELT-2 levels throughout the worm’s lifespan and ensure the intestine fate is maintained. Positive-feedback auto-regulation of this type has been shown to ensure the TF *che-1* can establish adequate expression intensity during differentiation and maintain expression throughout the worm’s lifespan to promote ASL neuronal identity (Leyva-Díaz and Hobert 2019). Indeed, heat shock-induced ectopic over-expression of ELT-2 expression leads to ectopic *elt-2* promoter reporter activity (Fukushige et al. 1998; Sommermann et al. 2010; Wiesenfahrt et al. 2015).

However, the ELT-2 positive autoregulation hypothesis had yet to be evaluated by removing ELT-2 from the biological system. We found that loss of ELT-2 led to an upregulation of *elt-2* promoter activity in the intestine. Additionally, we identified that simultaneous depletion of ELT-2 and ELT-7 reduced *elt-2* promoter activity back to wild-type levels (Figure 7). We propose two mechanisms for how ELT-2 and ELT-7 may regulate the *elt-2* promoter. ELT-2 and ELT-7 may both compete for the *elt-2* promoter which they can both activate, with ELT-7 serving as a stronger activator (Figure 8C). In this scenario, the balanced association of both ELT-2 and ELT-7 at the *elt-2* promoter creates a medium level of output, but removal of ELT-2 allows ELT-7 exclusive access thereby promoting a higher level of expression. Alternatively, ELT-2 may perform direct repression at the *elt-2* promoter by recruiting repressive machinery (Figure 8D). In this scenario, ELT-2 and ELT-7 create a “push and pull” of transcriptional output at the *elt-2* promoter. Finally, the observation that *elt-2* promoter activity is still expressed even in the absence of both ELT-2 and ELT-7 protein suggests an unknown TF likely also contributes to its activation.

We now have evidence that ELT-2 can both activate and repress its own promoter’s transcriptional output. ELT-2 can stimulate its promoter as shown by the fact that heat-shock-inducible ELT-2 stimulates expression of an *elt-2* promoter reporter (Sommermann et al. 2010). The evidence that ELT-2 can repress its own promoter comes from this study where we have shown that ELT-2 depletion leads to an over-expression of an *elt-2* promoter reporter. How can both scenarios be true? First, it is possible that the two experimental situations yield different results due to some subtleties of context. Second, it may indeed be possible that ELT-2 can act as both an auto-activator and auto-repressor through the direct recruitment of two classes of transcriptional regulatory machinery, perhaps directed by differences in ELT-2 concentration or developmental timing. Or third, ELT-2 may be competent to simultaneously activate and repress itself through the GRN network that may be organized into complex incoherent and/or paradoxical feedback loops. Indeed, the ELT-2 homolog ELT-7 is required for the repressive task suggesting that the GRN is involved.

We also determined that ELT-2 positively regulates some direct targets, negatively regulates others, and does not seem to transcriptionally impact the remainder of direct targets alone. Several examples of GATA factors in other organisms have revealed they are capable of both activation and repression through two mechanisms: competition or direct repression (Bresnick et al. 2010; Zheng and Blobel 2010; Block and Shapira 2015; Fujiwara 2017). Over-expression of human GATA factors GATA-4, GATA-5, and GATA-6 can inhibit tetracycline-catalyzed induction by out-competing higher activity tetracycline activators due to naturally occurring GATA sites within the tetO promoter (Gould and Chernajovsky 2004), illustrating a competitive mechanism of repression. In contrast, numerous examples of direct repression have been documented (Tremblay et al. 2018). In humans, GATA factors catalyze cell differentiation, act as either tumor suppressors or oncogenes, and serve as critical markers of cancer onset and progression (Fujikura et al. 2002; Bresnick et al. 2010; Chou et al. 2010; Zheng and Blobel 2010; Fujiwara 2017). Human GATA-1 positively regulates subsets of direct targets through the recruitment of histone acetyltransferase complexes (CBP/p300) (Blobel et al. 1998) but represses others through recruitment of direct repressive factors such as PU.1, FOG-1, or the NuRD co-repressor complex (Hong et al. 2005; Burda et al. 2010). In yeast, GATA factors have also shown to engage in direct repression. The yeast GATA factor Gaf1 responds to amino acid depletion by activating amino acid biosynthetic genes through RNA Pol II activation while simultaneously repressing tRNA transcription through RNA Pol III repression (Rodríguez-López et al. 2020).

Future experimentation will address the questions we have raised. A detailed dissection of the ELT-2 promoter could determine whether activating and repressing sequences can be isolated. However, because the binding sites for ELT-2, ELT-7 (and even PQM-1) are all identical, interpreting those experiments will be challenging. A dissection of the ELT-2 protein itself may be fruitful to determine whether it contains separate domains to recruit activating or repressing machinery. However, no such domains are predicted from the sequence. What will be necessary is an investigation into the inter-reliance between ELT-2, ELT-7, and PQM-1 – all TFs that bind identical sequences. And it is clear from our work that there are missing components of the intestinal GRN, particularly in the embryo, given that ELT-2 activity alone accounts for a minor fraction of intestines-specific gene expression at that stage. Overall, how tissue specific GRNs change dynamically over developmental time is poorly understood in any system. But, due to its past foundational work and overall tractability, the *C. elegans* intestinal GRN system is well poised to make inroads into this important field in future studies.

## Data availability

Strains and plasmids are available upon request. Supplemental material is available through figshare. Detailed protocols for FACS isolation of intestine cells are in the following protocols.io collection https://dx.doi.org/10.17504/protocols.io.5jyl895jdv2w/v1. Code for computational analysis is available in the GitHub repository https://github.com/rtpwilliams/williams_et_al. Sequencing data collected is available through NCBI Gene Expression Omnibus (GEO) database under accession number GSE214581 and accessible at https://www.ncbi.nlm.nih.gov/geo/query/acc.cgi?&acc=GSE214581. The gtf alignment files used in this analysis are also deposited there. The raw sequencing files are available at SRA (Short Read Archive) under Project PRJNA885900.

## Acknowledgments

We thank Dr. James McGhee, Michelle Kudron, Valerie Reinke, and WormBase for protocols, equipment, reagents, strains, advice, productive discussions, feedback regarding the manuscript, and commentary. Some strains were provided by the CGC, which is funded by NIH Office of Research Infrastructure Programs (P40 OD010440). This work utilized resources from the University of Colorado Boulder Research Computing Group, which is supported by the National Science Foundation (awards ACI-1532235 and ACI-1532236), the University of Colorado Boulder, and Colorado State University. This work utilized microscopy resources from NIH grant 1S10 OD025127 and support from the CSU Microscope Imaging Network. Some Figure elements were created in BioRender.

