## supplemental for "Transcriptome profiling of the *Caenorhabditis elegans* intestine reveals that ELT-2 negatively and positively regulates intestinal gene expression within the context of a gene regulatory network"

Genome-wide characterization of the *Caenorhabditis elegans* intestine GATA transcription factor ELT-2

#### Supplemental Figures and Legends:

**Figure S1. FACS gating strategy for isolation of intestine cells.** Diagram of gating strategy used for embryo (A), L1 (B), and L3 (C) stage FACS intestine isolation.

**Figure S2. Sample-to-sample Euclidean distances of embryo and L1 stage RNA-seq samples.** Pairwise measurements for Euclidian distance of regularized logarithm transformed RNA-seq read counts between intestine and non-intestine RNA-seq libraries for assayed developmental stages. L1 stage samples from replicate 2 clustered with embryo stage samples and were removed from downstream RNA-seq analysis.

**Figure S3**. **Identification of dissociation-induced transcript abundance difference in C. elegans single cell suspension compared to whole worms. (A)** MA plots visualizing differential transcript abundance is dissociated cells compared to whole worms. Points are colored based on the category: significantly unchanged (blue), reduced in dissociated cells (green), increased in dissociated cells (red). **(B)** Bar plot displaying the number of transcripts in each category.

**Figure S4. ELT-2 binding status is associated with higher mean expression of intestine-enriched genes in every stage.** Violin plot of genes intestine expression for genes categorized based on presence or absence of ELT-2 peak in target gene promoter. The number of genes in each set are indicated below each histogram. Student's t-test was used to measure statistical significance. Bars within the violin plot represent the 99% confidence interval.

**Figure S5. Intestine expression is not correlated to ELT-2 occupancy of target gene promoters. (A)** As a quantitative predictor, the relationship between ELT-2 occupancy signal and gene expression is not higher for genes associated with a promoter that is reproducibly bound by ELT-2 (slopes are the same or lower in magnitude between the two binding statuses, for each stage independently). The greater $R^2$ values for ELT-2 bound data indicate that those data fit their respective models better, but still reflect a relatively low variance explained (.10 - .14). **(B)** Slopes did not improve for when genes were further subset based on dependence for ELT-2 transcriptional regulation. Thin black lines represent contour lines for point density.

**Figure S6. Visualization of ELT-2 ChIP-seq read density averaged across promoter regions centered on the gene transcription start site.** Promoters were defined as -1kb upstream and 0.2kb downstream of a gene transcription start site (dark blue, ELT-2 bound and intestine enriched; light blue, ELT-2 bound and not intestine enriched; ELT-2 not bound, intestine enriched; ELT-2 not bound, not intestine enriched).

**Figure S7. Visualization of repressed ELT-2 target gene locus. (A-C)** Genome browser tracks of the *cebp-1* (A), *ets-4* (B), and *pqm-1* (C) genomic locus. RNA-seq tracks (blue) display increased transcript abundance in *elt-2(-)* genetic background compared to wildtype (WT, N2). ELT-2 ChIP-seq tracks (black) display ELT-2 binding sites in the gene promoters (black bars).

**Figure S8. Negative regulation of the *elt-2* promoter is observed in L1, L3 and adult stages. (A)** Block diagram of the *elt-2* promoter reporter allele used in this study. **(B)** Quantification of elt-2 promoter intestine fluorescence in either control RNAi (L4440) or *elt-2* RNAi conditions. Measurements were performed in L1, L3 and adult stages. GFP measurements were collected from background subtracted maximum Z projections from three biological replicates and normalized for intestine area (n = 30 worms for all stages and RNAi treatments). Points represent a single worm and are colored based on biological replicate. Data distribution is represented by box and whisker plots. Student's t-test was used for significance testing. Plot y-axis is log-scale. **(C)** Relative fluorescence of *elt-2* promoter reporter activity measured in *elt-2* RNAi over control RNAi in L1, L3 and adult developmental stages. Error bars represent the t-test 95% confidence interval for the ratio of means between *elt-2* RNAi and control RNAi measurements.

**Figure S9. Visualization of ELT-2 negative autoregulation in RNA-seq data. (A)** Genome browser tracks of the *elt-2* locus. RNA-seq tracks (blue) display increased read density aligning to the *elt-2* promoter in *elt-2(-)* genetic background compared to wildtype (WT, N2). The increase in *elt-2* promoter read density is reduced in *elt-2(-);elt-7(-)* genetic background. **(B)** Quantification of RNA-seq reads aligning to the elt-2 promoter. The elt-2 promoter here is defined as the region spanning the TSS to -1kb upstream of the TSS. RNA-seq reads were normalized with the DESeq2 package.

**<u>Supplemental Files:</u>**

**File S1:** Table of samples collected for FACS-isolated intestine RNA-seq.

**File S2:** Raw, normalized and, regularized log (rlog) transcript count tables for FACS-isolated intestine RNA-seq data.

**File S3:** Pairwise results tables for intestine (GFP+) vs. non-intestine (GFP-) samples for embryo, L1 and L3 stages.

**File S4:** ELT-2 ChIP-seq data files utilized from the modERN Resource.

**File S5:** Data tables containing the integration of ELT-2 ChIP-seq, elt-2 (-) RNA-seq and FACS-isolated intestine RNA-seq data in embryo, L1 and L3 stages.

**File S6:** Non-overlapping anatomy ontology terms (See Methods, "Gene set definitions").

**File S7:** Regression statistics for correlation of ELT-2 ChIP-seq promoter occupancy and intestine transcript abundance.

**File S8:** smiFISH probe sequences utilized in this study.

Figure S1

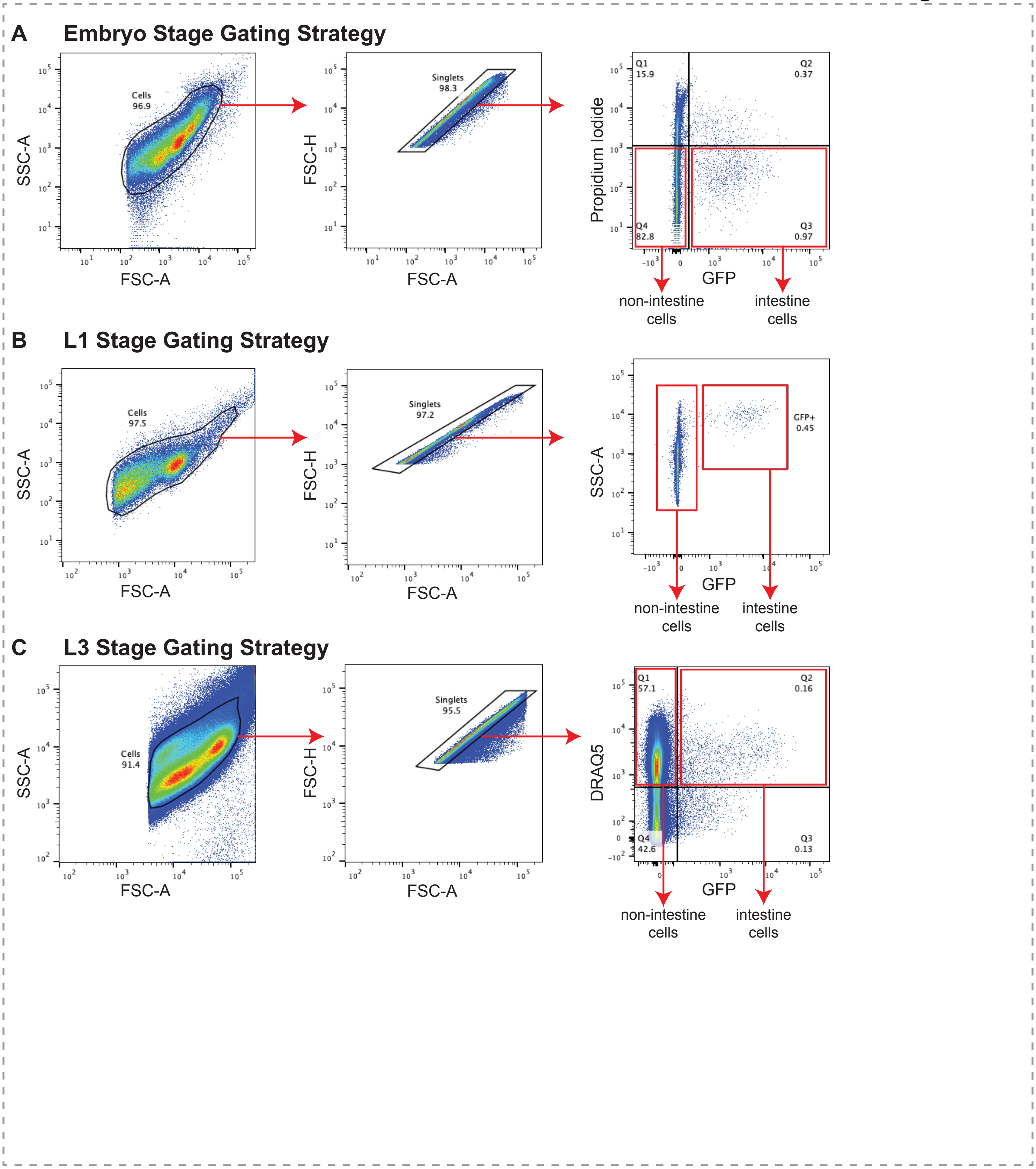

A
Embryo Stage Gating Strategy
Cells
96.9
SSC-A
FSC-A
Singlets
98.3
FSC-H
FSC-A
Q1
15.9
Q2
0.37
Q4
82.8
Q3
0.97
Propidium Iodide
GFP
non-intestine cells
intestine cells
B
L1 Stage Gating Strategy
Cells
97.5
SSC-A
FSC-A
Singlets
97.2
FSC-H
FSC-A
GFP+
0.45
SSC-A
GFP
non-intestine cells
intestine cells
C
L3 Stage Gating Strategy
Cells
91.4
SSC-A
FSC-A
Singlets
95.5
FSC-H
FSC-A
Q1
57.1
Q2
0.16
Q4
42.6
Q3
0.13
DRAQ5
GFP
non-intestine cells
intestine cells

Figure S2

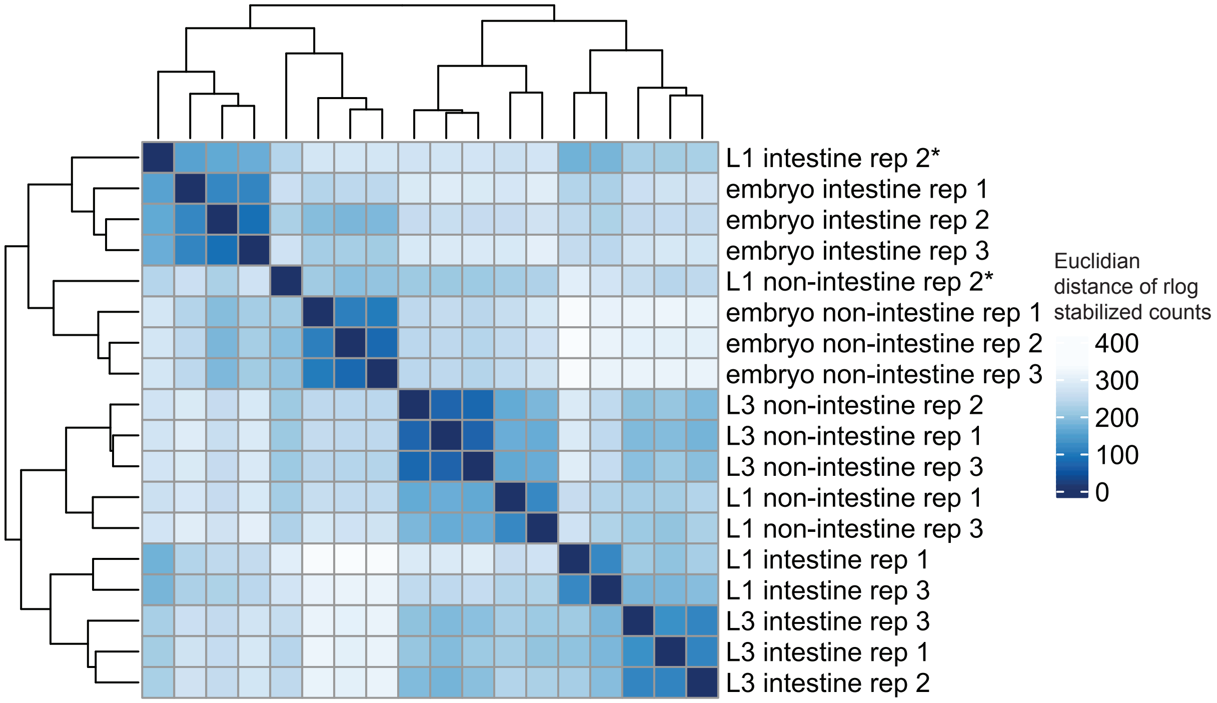


*samples removed from downstream analysis

### Figure S3

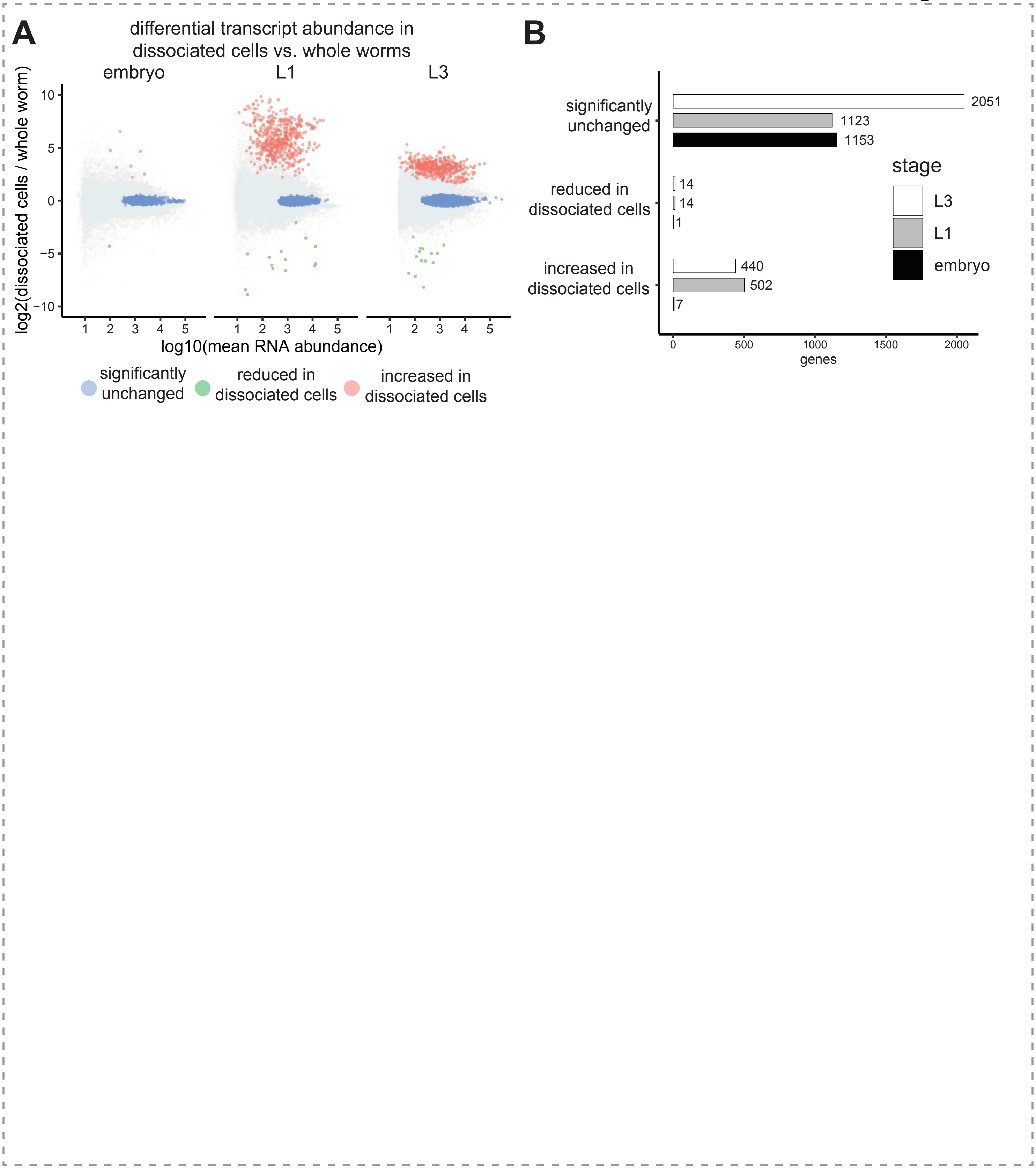

A
differential transcript abundance in dissociated cells vs. whole worms
embryo
L1
L3
log2(dissociated cells / whole worm)
10
5
0
−5
−10
1 2 3 4 5
log10(mean RNA abundance)
significantly unchanged
reduced in dissociated cells
increased in dissociated cells
B
significantly unchanged
2051
1123
1153
reduced in dissociated cells
14
14
1
increased in dissociated cells
440
502
7
0 500 1000 1500 2000
genes
stage
L3
L1
embryo

### Figure S4

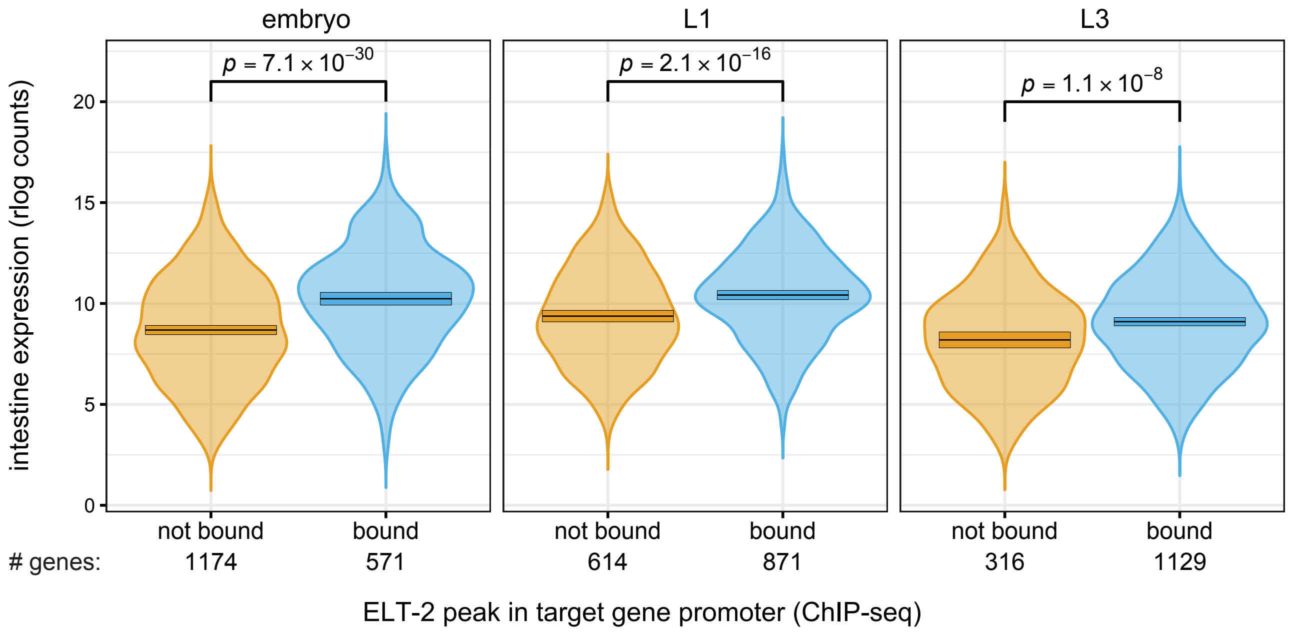

embryo
L1
L3
$p = 7.1 \times 10^{-30}$
$p = 2.1 \times 10^{-16}$
$p = 1.1 \times 10^{-8}$
intestine expression (rlog counts)
0
5
10
15
20
not bound
bound
### genes:
1174
571
614
871
316
1129
ELT-2 peak in target gene promoter (ChIP-seq)

### Figure S5

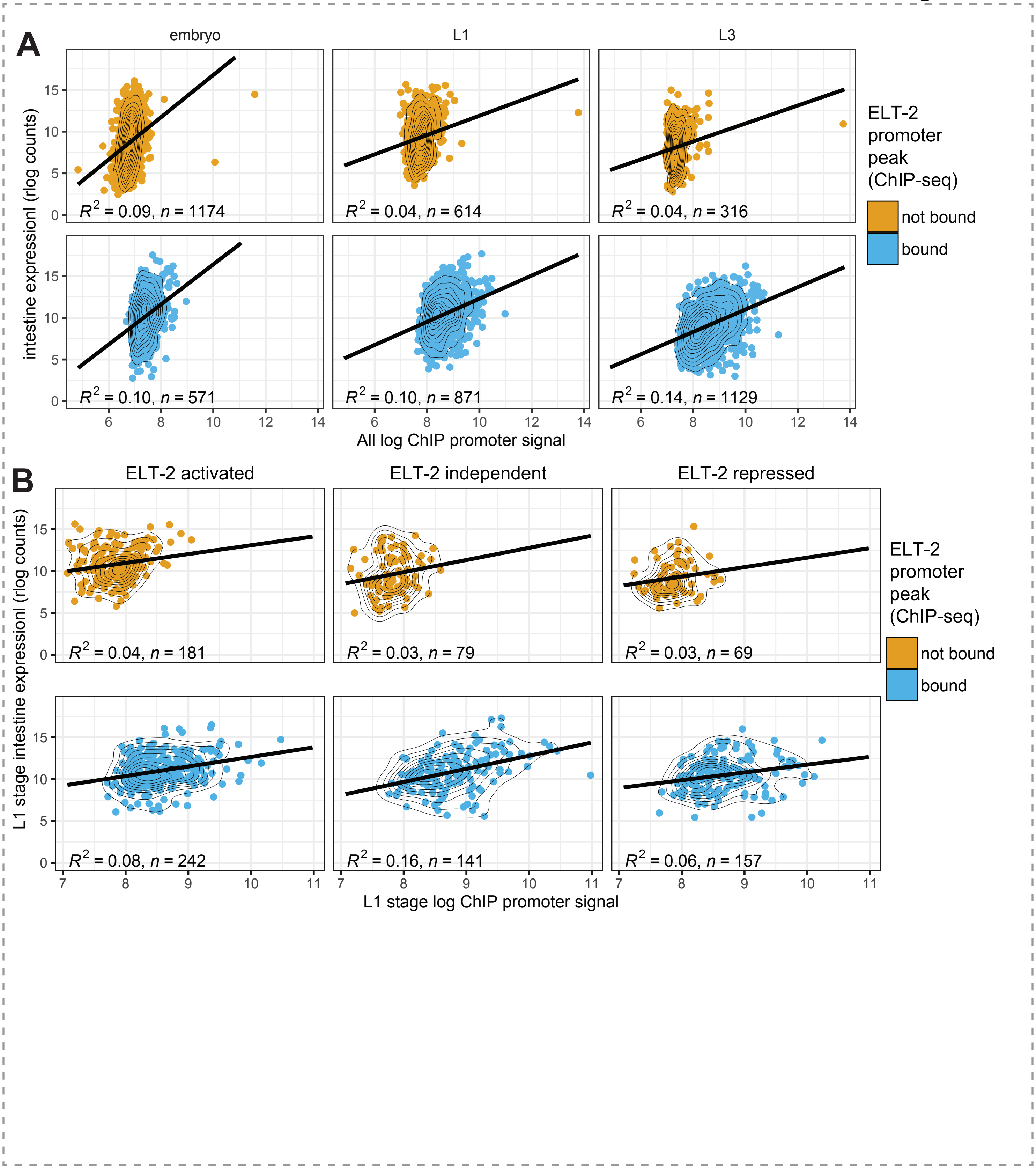
A
embryo
L1
L3
intestine expressionl (rlog counts)
$R^2$ = 0.09, $n$ = 1174
$R^2$ = 0.04, $n$ = 614
$R^2$ = 0.04, $n$ = 316
$R^2$ = 0.10, $n$ = 571
$R^2$ = 0.10, $n$ = 871
$R^2$ = 0.14, $n$ = 1129
All log ChIP promoter signal
ELT-2 promoter peak (ChIP-seq)
not bound
bound
B
ELT-2 activated
ELT-2 independent
ELT-2 repressed
L1 stage intestine expressionl (rlog counts)
$R^2$ = 0.04, $n$ = 181
$R^2$ = 0.03, $n$ = 79
$R^2$ = 0.03, $n$ = 69
$R^2$ = 0.08, $n$ = 242
$R^2$ = 0.16, $n$ = 141
$R^2$ = 0.06, $n$ = 157
L1 stage log ChIP promoter signal
ELT-2 promoter peak (ChIP-seq)
not bound
bound

### Figure S6

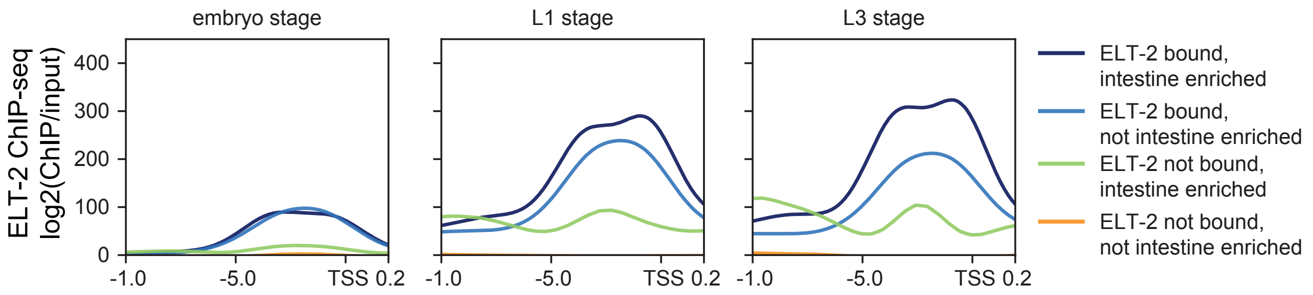
embryo stage
L1 stage
L3 stage
ELT-2 ChIP-seq
log2(ChIP/input)
400
300
200
100
0
-1.0
-5.0
TSS 0.2
ELT-2 bound, intestine enriched
ELT-2 bound, not intestine enriched
ELT-2 not bound, intestine enriched
ELT-2 not bound, not intestine enriched

Figure S7

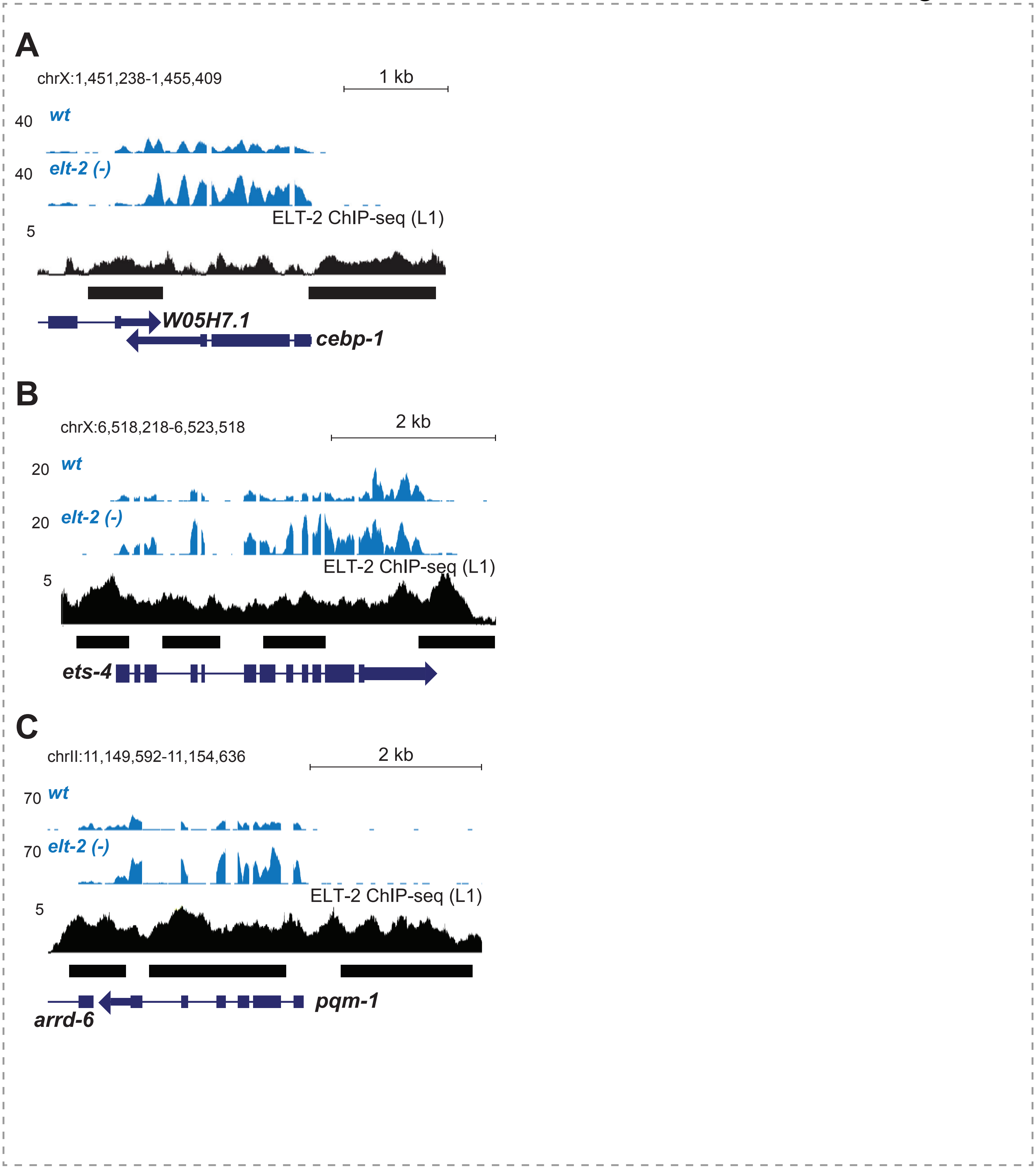
A
chrX:1,451,238-1,455,409
1 kb
40
wt
40
elt-2 (-)
ELT-2 ChIP-seq (L1)
5
W05H7.1
cebp-1
B
chrX:6,518,218-6,523,518
2 kb
20
wt
20
elt-2 (-)
ELT-2 ChIP-seq (L1)
5
ets-4
C
chrII:11,149,592-11,154,636
2 kb
70
wt
70
elt-2 (-)
ELT-2 ChIP-seq (L1)
5
pqm-1
arrd-6

Figure S8

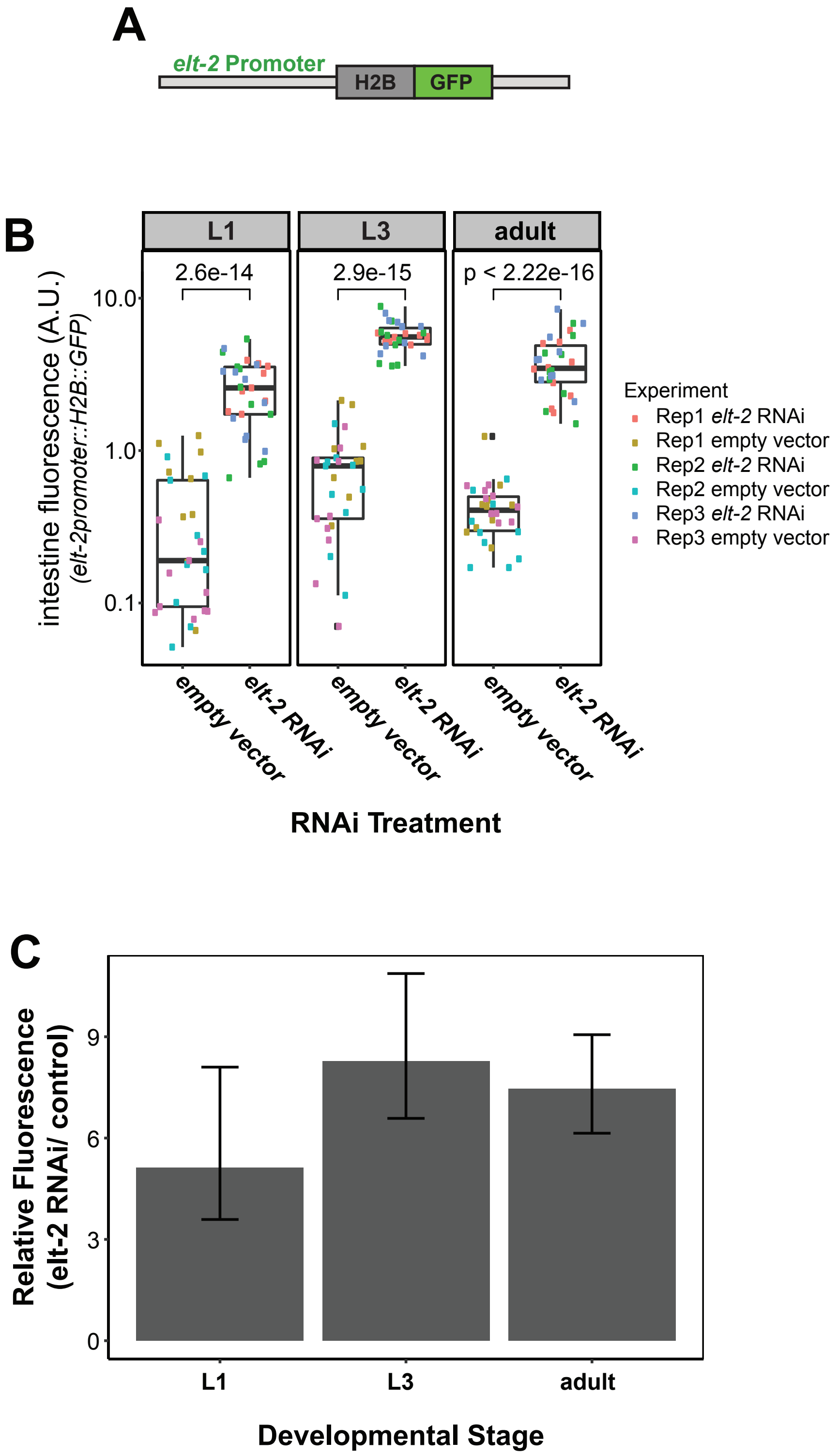
A
elt-2 Promoter
H2B
GFP
B
L1
L3
adult
2.6e-14
2.9e-15
p < 2.22e-16
intestine fluorescence (A.U.)
(elt-2promoter::H2B::GFP)
10.0
1.0
0.1
empty vector
elt-2 RNAi
RNAi Treatment
Experiment
Rep1 elt-2 RNAi
Rep1 empty vector
Rep2 elt-2 RNAi
Rep2 empty vector
Rep3 elt-2 RNAi
Rep3 empty vector
C
Relative Fluorescence
(elt-2 RNAi/ control)
9
6
3
0
L1
L3
adult
Developmental Stage

Figure S9

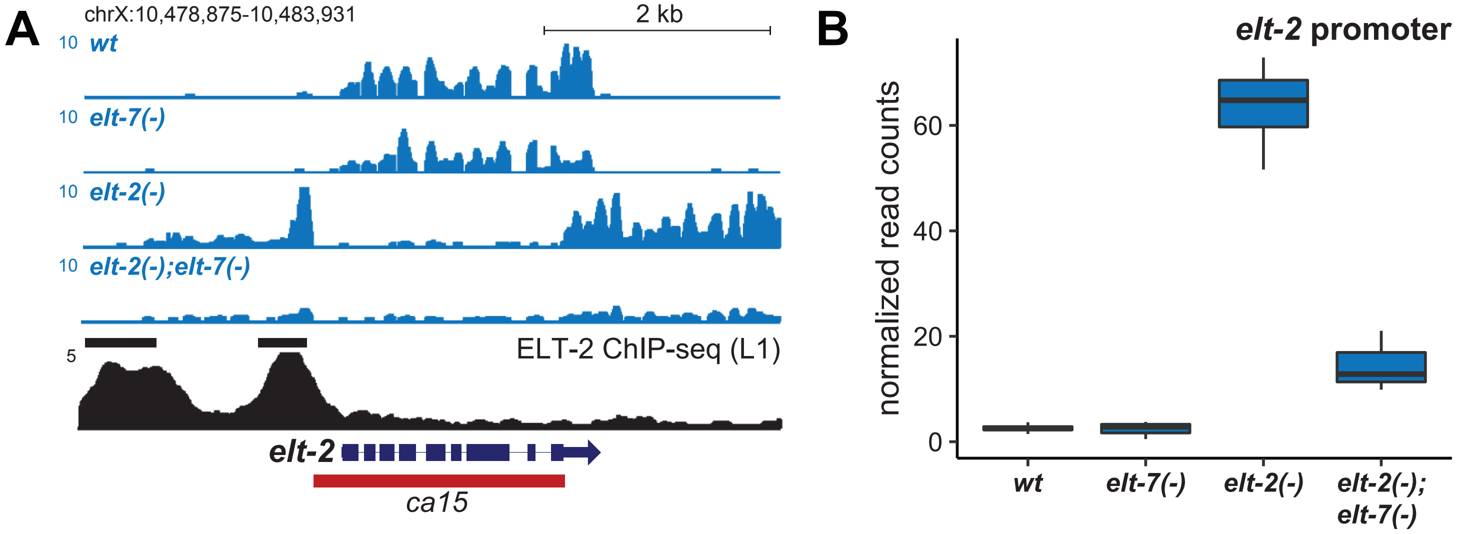

A
chrX:10,478,875-10,483,931
2 kb
10 wt
10 elt-7(-)
10 elt-2(-)
10 elt-2(-);elt-7(-)
5
ELT-2 ChIP-seq (L1)
elt-2
ca15
B
elt-2 promoter
normalized read counts
60
40
20
0
wt
elt-7(-)
elt-2(-)
elt-2(-);
elt-7(-)
